# A Biophysical Model for Plant Cell Plate Development

**DOI:** 10.1101/2020.05.21.109512

**Authors:** Muhammad Zaki Jawaid, Rosalie Sinclair, Daniel Cox, Georgia Drakakaki

**Affiliations:** Department of Physics, University of California, Davis, 95616; Department of Plant Sciences, University of California, Davis, 95616

**Keywords:** Cell Plate, Cytokinesis, Callose, Spreading Force, Helfrich Energy

## Abstract

Plant cytokinesis, a fundamental process of plant life, involves *de novo* formation of a ‘cell plate’ that partitions the cytoplasm of the dividing cell. Cell plate formation is directed by orchestrated delivery, fusion of cytokinetic vesicles, and membrane maturation to the form the nascent cell wall by the timely deposition of polysaccharides such as callose, cellulose, and crosslinking glycans. In contrast to the role of endomembrane protein regulators the role of polysaccharides, in cell plate development is poorly understood. Callose, a β-1-3 glucan polymer, is transiently accumulated during cell plate expansion to be replaced by cellulose in mature stages. Based on the severity of cytokinesis defects in the absence of callose, it has been proposed that it stabilizes this membrane network structure. However, there is currently no theory to understand its role in cytokinesis.

Here we extend the Helfrich free energy model for membranes including a phenomenological spreading force as an “areal pressure” generated by callose and/or other polysaccharides. Regular cell plate development in the model is possible, with suitable bending modulus, for a two-dimensional late stage spreading force parameter of between 2–6*pN*/*nm*, an osmotic pressure difference of 2–10*kPa*, and spontaneous curvature between 0–0.04*nm*^−1^. With these conditions, stable membrane conformation sizes and morphologies emerge in concordance with stages of cell plate development. With no spreading force, the cell plate fails to mature properly, corroborating experimental observations of cytokinesis arrest in the absence of callose. To reach a nearly mature cell plate, our model requires the late stage onset that the spreading force coupled with a concurrent loss of spontaneous curvature. A simple model based upon production of callose as a quasi-two-dimensional self-avoiding polymer produces the correct phenomenological form of the spreading force, which will be further refined, since matching to our numbers requires an exceptionally high callose synthesis rate.

**Significance Statement:** Plant cell division features the development of a unique membrane network called the cell plate that matures to a cell wall which separates the two daughter cells. During cell plate development, callose, a β-1-3 glucan polymer, is transiently synthesized at the cell plate only to be replaced by cellulose in mature stages. The role for this transient callose accumulation at the cell plate is unknown. It has been suggested that callose provides mechanical stability, as well as a spreading force that widens and expands tubular and fenestrated cell plate structures to aid the maturation of the cell plate. Chemical inhibition of callose deposition results in the failure of cell plate development supporting this hypothesis. This publication establishes the need for a spreading force in cell plate development using a biophysical model that predicts cell plate development in the presence and the absence of this force. Such models can potentially be used to decipher for the transition/maturation of membrane networks upon the deposition of polysaccharide polymers.

## Introduction

A unique compartment of plant cells is their cell wall, a dynamic network of polysaccharides and other molecules that provides both structural support and protection from external stresses (1). The plant cell wall mechanical heterogeneities are able to regulate cell asymmetry and morphogenesis, a research area that has received much attention recently (2). While many studies focus on the rigidity of cell wall to constrict shape against the turgor pressure, little is known on polysaccharide driven membrane network expansion.

De novo formation of cell walls occurs during cytokinesis in plants. In plants, formation of a cell plate develops into the new cell wall, partitioning the cytoplasm of the dividing cell. Cell plate formation takes place in multiple stages that involve highly orchestrated vesicle accumulation, fusion and membrane transformation concurrent with the time specific deposition of polysaccharides such as callose, cellulose and cross-linking glycans. This development requires the directed and choreographed accumulation of post-Golgi vesicles via the phragmoplast, which is an assembly of microtubules and microfilaments that helps organize vesicle delivery to the cell plate assembly matrix, at the division plane.

Cell plate expansion is centrifugal, led by the accumulation and fusion of cytokinetic vesicles to the leading edge (3-6). The different cell plate development stages are defined morphologically as follows (Fig. 1). During the initial fusion of Golgi vesicles stage (FVS), cytokinetic vesicles, mainly derived from the Golgi, are guided by the phragmoplast to the cell plate assembly matrix where their fusion starts to occur (Fig. 1*A*). Fused vesicles are transformed into dumbbells that undergo conformational changes to form a tubulo-vesicular network (TVN) (Fig. 1*B*), which then expands to the tubular network (TN) (Fig. 1*C*). The TN then expands into a planar fenestrated sheet that ultimately develops into a young cell wall (3, 4). Excess membrane material during the transitional stages is recycled concurrently, along with the accumulation of different polysaccharide materials (4). It is important to note that multiple stages exist simultaneously, adding complexity in dissecting them (7).

**Figure 1.**
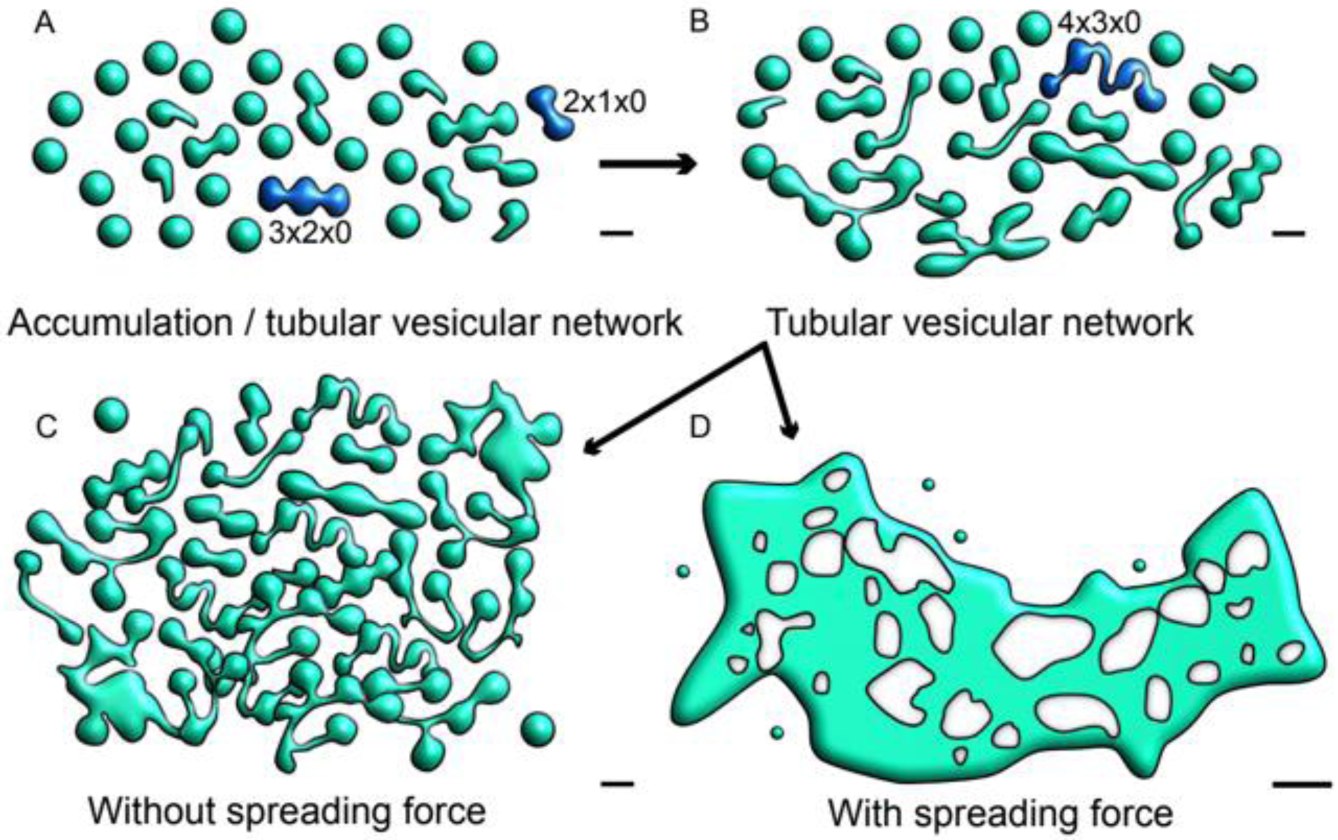
Schematic representation of cell plate development describing the role of a spreading force. Early stages of vesicle accumulation and fusion (*A*), and tubular network structures are shown (*B*). Two different possibilities are projected for stage transition. I) Incomplete/arrested cell plate (*C*). In the absence of a spreading force (*C*), tubular and fenestrated structures are accumulating, and there is a lack of maturation towards a single, complete cell plate structure. II) Normal cell plate transition (*D*). In our calculations, we discover that for expansion/maturation to occur as in (*D*), the presence of a spreading force is required, along with the decrease of spontaneous curvature to a threshold value. This allows for a sheet like cell plate structure (SCP) to form. The structures in this schematic description are adapted from data collected from EM tomography (2) with bars in *A* to *C* = 50nm, *D* = 0.25µm. Dark blue vesicles denote those labeled by the mathematical naming schema as described in Fig. 2. Where in A, 2×1×0 denotes 2 oblate spheroids, 1 tubular connection and 0 holes.

Although several determinants involved in cell plate formation have been identified (5), a biophysical understanding of the contribution of the molecular and structural players involved in cell plate expansion is not available. During cell plate expansion, membrane remodeling and network expansion is highly coordinated with the deposition of polysaccharides, which therefore, provides an excellent opportunity to study membrane morphology changes introduced by polymer deposition. Currently, most of the biophysical models are concentrated in the role of curvature-stabilizing proteins to induce membrane morphology changes. These include curvature shaping proteins in cross-sections of tubules and sheet edges of endoplasmic reticulum (8-10) among an array of studies in diverse systems. Furthermore, the current models focus on the protein activity while limited information exists on membrane expansion driven by polysaccharide polymer generation.

The development of new cell walls during cell division is of profound importance in all plant life, yet fundamental questions about this process remain unanswered, including: How does callose contribute to cell plate expansion? Callose, a β-1-3 glucan polymer, is transiently synthesized at the cell plate and its accumulation peaks at the intermediate TN stage. Currently, it is not known why callose is deposited in the developing cell plate during intermediate stages (namely, the TVN, TN and PFS), only to be replaced by cellulose (a β-1-4 glucan polymer) close to maturation to a complete cell wall (3).

It is widely believed that callose provides mechanical stability in addition to a rapid transient spreading force that helps transition the membrane network to a more developed stage (3) (Fig. S1). However, there is no biophysical understanding of whether a spreading force is necessary for cell plate expansion in the first place.

Studies have revealed that this energetically costly temporal inclusion of callose is vital to cell plate development (11, 12). Genetic mutations in the cytokinesis specific callose synthase (glucan synthase like 8, GSL8) are lethal (11, 13, 14). In addition, selective chemical inhibition of callose synthase by the small molecule Endosin 7 (ES7) arrests late stage cytokinesis with a typical bi-nucleate cell phenotype (12). The lethality induced by callose inhibition in cytokinesis, in combination with the recalcitrance of the recombinant GSL8 callose synthase, provides significant hurdles to detailed studies on the regulation of callose synthase activity.

We hypothesize that callose provides a necessary time-dependent spreading force, which we can model by adding a phenomenological “areal pressure” term to the Helfrich model free energy for the cell plate surface, and we can study the influence of this spreading force by adopting a variational approach to locally minimize the model free energy in time, assuming the process is sufficiently slow that we are close to thermodynamic equilibrium. The quasi-equilibrium is constantly redefined as vesicles are added at the cell plate boundary. This enables us to use the total cell plate surface area as a proxy for time. We demonstrate semi-quantitatively that by assuming a late time onset of the spreading force followed by the reduction of membrane spontaneous curvature, that we can reproduce the observed morphological time course of the cell plate. We do not have a detailed microscopic model for the spreading force, but we do show that a simple model based upon the expansion of callose as a quasi-two-dimensional self-avoiding polymer captures the correct form. A detailed effort to match the model to the estimated spreading force coefficient from the free energy requires excessively high callose production rates.

## Materials and Methods

### Energy Minimization

For our energy minimization calculations, we treat the membrane boundary of the cell plate as an incompressible two-dimensional surface to a first approximation, with certain defined characteristics. In reality, each lipid bilayer has a finite thickness of 4-6nm (15), however, it is justified to treat the surface of the cell plate as two-dimensional due to the smallness of its thickness compared to the sizes of the overall structures in the cell plate. It can also be shown that lipid bilayers present a high level of incompressibility due to the energy penalty associated with areal stretching being significantly higher compared to membrane deformations due to bending (16). This is similar to the approach taken by Choksi et al.(17) and Sarasij et al.(18).

At a specified area, we define a free energy of our membrane surface that is essentially the Helfrich energy (19) with the addition of a novel term to model the spreading force, possibly due to callose, as given in Eq. (1). We discuss the possible origin of this spreading force within a simple model elsewhere. Finally, we consider the surface area of the cell plate as a proxy for cell plate development time and then we minimize this energy for a given surface area.

Due to the complexity of this research problem, we decided to look for energy minima within a parameterized restrictive geometry basis set, thereby adopting a restricted variational approach. We found that existing adaptive mesh approaches such as Surface Evolver, while in principle more accurate and flexible, were not readily amenable for application in our study (20).

### Shape Approximation

Structures that are found within the vesicular cloud can be approximated using oblate spheroids, where the two defining radii can be used as variational parameters as shown in Fig. 2*A*. The oblate spheroid can also be used to model the expanded/late stage cell plate close to completion, as a very large oblate spheroid with *a* ≫ *c*.

**Figure 2.**
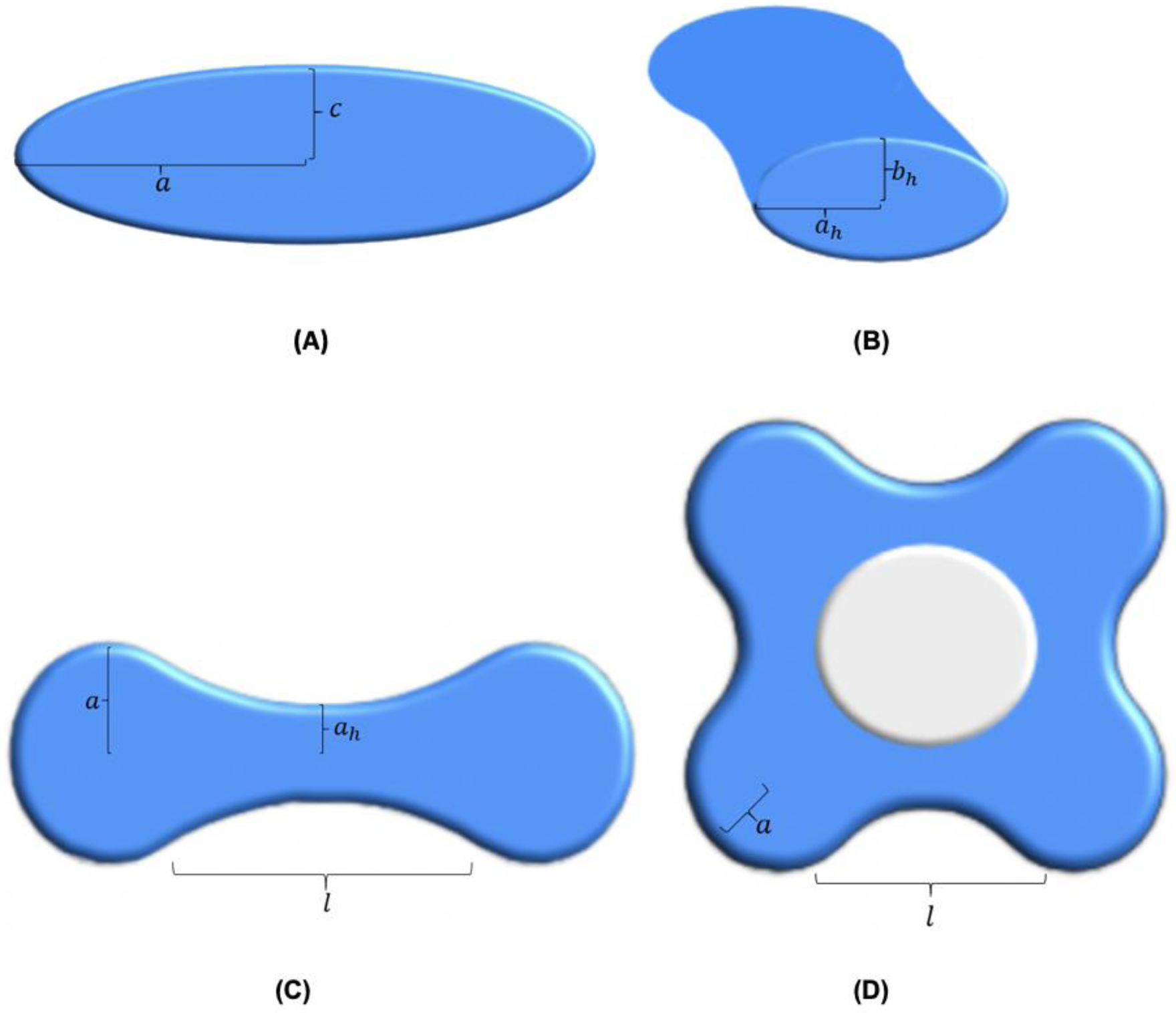
Examples of membrane structure parameterizations used for modeling. (*A*) Cross section of an oblate spheroid through the polar axis. The major axis radius is labelled *a* and the minor axis radius is labelled *c*. This structure is used to model vesicles, or larger cell plate structures in the case where *a* ≫ *c*. (*B*) A cross section of an elliptic hyperboloid at its center, showing the skirt radii. The hyperboloid can be parameterized by its length *l* and its skirt radius in the equatorial plane *a*_*h*_, the skirt radius in the axial plane is given by *b*_*h*_, which can be written as a function of the other parameters listed as shown in Eq. (S3) (*C*) An example of a tubulo-vesicular structure parameterized by two oblate spheroids connected by a single elliptic hyperboloid (referred to as a 2×1×0 structure). Only the top view is shown. (*D*) An example of a 4×4×1 conformation that models a transition to a fenestrated network with genus *g* = 1.

Similarly, structures found within the fenestrated sheet and the tubulo-vesicular network stages can be approximated using a combination of elliptic hyperboloids and oblate spheroids, such that the hyperboloids are continuous at the oblate spheroid boundaries. These elliptic hyperboloids can be parameterized by their length, hereafter referred to as *l*, and their skirt radius in the equatorial plane, hereafter referred to as *a*_*h*_. The other parameters needed to define an elliptic hyperboloid can be written as a function of the two parameters already listed due to boundary conditions that arise from mandating continuity. Fig. 2*B* shows a cross section of an elliptic hyperboloid, while Fig. 2*C* shows an example of two vesicles joined by a single tube.

### Naming convention of different approximated conformations

The naming convention for different conformations was defined as described in the following example: A conformation that is labeled as 6×7×2, represents a conformation that has 6 oblate spheroids (or vesicles), 7 hyperboloids (or tubes), and 2 holes (or fenestrations, with *g* = 2). A 2×1×0 conformation is shown in Fig. *2C*, and an example of a 4×4×1 conformation with a single fenestration is shown in Fig. *2D*.

The area of any given conformation was calculated by numerical integration methods, and a corresponding parameter space of a given area was found. For simplicity, we analyzed conformations that shared the same parameters for each of the oblate spheroids, and each of the hyperboloids with examples shown in Fig. *2* with additional examples in Fig. 1. Therefore, we found a four-dimensional parameter space (*a, c, a*_*h*_, *l*) that corresponded to a given area up to an error tolerance (<0.05%) for each conformation of interest, and then calculated the energies of Eq. (1) within that parameter space. The energy minimum was then extrapolated from that parameter space. Additional information about extrapolating a four-dimensional parameter space is given in the supplemental information (Eq. (S1-S4)), and a full list of the parameters involved are shown in Fig. S2.

### Plant Growth

Arabidopsis seedlings of Col-0 were used in this study. Seeds were sterilized using 30% (v/v) sodium chlorate in ethanol (absolute) with 0.06% (v/v) of Triton X-100 (Sigma-Aldrich). Seeds were plated on 0.25 Murashige and Skoog medium (1.15 g L^−1^ Murashige and Skoog minimal organics salt, 10 g L^−1^ Suc, 5 g L^−1^ Phytagel (Sigma-Aldrich), and cold vernalized for 48 h at 4°C in the dark, after which plates were transferred to a plant growth chamber for seedling growth. Plants were grown in temperature- and photoperiod-controlled environments, set to long-day (16-h-light/8-h-dark cycle) conditions, using fluorescent light (at 100 to 150 mmol quanta photosynthetically active radiation (PAR) m^−2^ s^−1^) at 22 to 24°C.

### Chemical treatment and Imaging

Four day old Arabidopsis seedlings were treated with 25 µM ES7 and DMSO in 0.25 MS medium for two hours as previously described in Ref. (12).

A Leica SP8 confocal microscope was used for imaging. Fluorescence signals of, Callose stained by Aniline blue fluorochrome (Biosupplies Australia) (excitation 350 nm, emission 450-510 nm), and FM4-64 (ThermoFisher Scientific) (excitation 488 nm, emission 652-759) were collected with 40x (water), 63x (oil) and 100x (oil) objectives. Z stacks were generated across the volume of full cell and were subsequently deconvolved with Huygens (SVI). 3D reconstructions were prepared using Imaris, Bitplane and figures were assembled using Adobe Illustrator.

## Results

In order to establish the role of callose and other polysaccharides in cell plate expansion we modeled the free energy for the cell plate surface by adopting the Helfrich energy (19) with the addition of a novel term to model the presence of a spreading force. The free energy is defined as follows: 

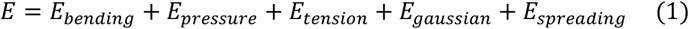

The first term describes the bending energy over the closed membrane surface(s) of the cell plate. It is given by: 

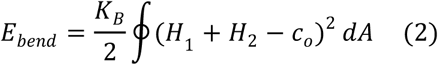

Where *K*_*B*_ is the bending modulus for the membrane surface, *H*_1_ and *H*_2_ are the principle curvatures at a point on the surface, and *c*_*o*_ is the spontaneous curvature, or the preferred curvature for the membrane. For simplicity, we assume that the bending modulus is time independent. We allow for the spontaneous curvature, *c*_*o*_, to be time dependent, which reflects potential differences in membrane composition during cell plate evolution.

The next term is the pressure energy which results from the difference in osmotic pressure between the inside and the outside of the cell plate, such that Δ*p* = *p*_*out*_ − *p*_*in*_. It is given by: 

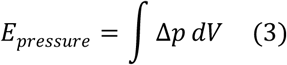

The third term is the energy associated with the surface tension of the membrane, given by: 

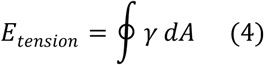

We assume that the surface tension given by *γ* is a constant. Since we assume that our system locally equilibrates in time at a constant area, this term only adds a constant energy equal to the area at that time, multiplied with the surface tension.

Finally, we introduce the novel element of a spreading force, which is analogous to a two-dimensional pressure that acts against the periphery of the cell plate structure along the equatorial plane. In this interpretation, *λ* has units of force/length, with the energy representation as follows: 

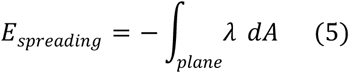

It is important to note that the integral in Eq.(5) is over the equatorial plane of closed cell plate surfaces. Therefore, fenestrations are not integrated over.

As with the spontaneous curvature, we allow for the spreading force coefficient *λ* to be time dependent, representing, e.g., the “turning on” of callose production in an expanding plate.

The last term is the Gaussian bending energy term, given by: 

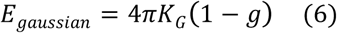

Here, *K*_*G*_ is the Gaussian bending modulus, and *g* is the genus of the surface. This is a result of the Gauss-Bonnet theorem(21). Finally, we consider the surface area of the cell plate as a proxy for cell plate development time and then we minimize this energy for a given surface area.

We use established values for the bending modulus *K*_*B*_ (22), as well as the Gaussian bending modulus *K*_*G*_ (23), and the surface tension *γ* (24). However, the pressure difference Δ*p* was phenomenologically determined. The bending modulus is sensitive to the environment of the cell as well as the membrane type and conformation (22), and therefore there is a range of available literature values. We decided to test over the full range of literature values ranging from about 62.5pNnm to 200pNnm (roughly corresponding to a range of 15*k*_*b*_*T* to 50*k*_*b*_*T*).

As described in the materials and methods section, we can model cell plate structures as combinations of oblate spheroids and elliptic hyperboloids. Oblate spheroids can be used to model vesicles, as well as an almost complete cell plate (in the case that *a* ≫ *c*), while combinations of oblate spheroids and elliptical hyperboloids can be used to model various tubular and fenestrated structures as shown in Figs. 1-2, S2. We identify our structures using a naming convention of (A_n_xB_n_xC_n_), where A_n_ is the number of oblate spheroids (or vesicles), B_n_ the number of hyperboloids (or tubes) and C_n_ the number of fenestrations (or perforations).

This methodology can also be used to provide a basis for the quantitative assessment of membrane structures found in the endoplasmic reticulum and the Golgi apparatus. There have of course been theoretical efforts to apply Helfrich theory to these structures, but few have taken a large scale view of the system. Notably, the work of Shemesh *et al.* (9) examines the morphologies possible as a function of curvature modifying proteins using full minimization of the free energy via the Surface Evolver finite element approach(20). The finite element methods are powerful and somewhat customizable, but we were unable to find any way to include the spreading force term together with the pressure energy term of Eq. (5) in any such code. In order to make progress, we adopted the variational approach including multiply connected surfaces with negative curvature tubulations as a reasonable compromise approach to explore the quasi-equilibrium stabilities of different morphologies that are fully representative of the observed structures.

Within this approach, we first calculated results for multiple types of structures to determine a range of parameters that would yield structures matching the experimentally observed cell plate sizes/thicknesses. From electron tomography Cryo-EM images of a developing cell plate, we determined that the thickness of a cell plate in various stages of development, with or without the presence of callose, was approximately 40-120nm(4). Therefore, we tuned the free parameters in our energy model such that the thickness of any of our conformations across the equatorial plane was in the range of 40-120nm. We determined that, depending on the choice of the bending modulus, the allowed values of the spreading force parameter *λ* should be between 0.0-6.0pN/nm, the spontaneous curvature *c*_*o*_ between 0-0.04*nm*^−1^, and a finite pressure difference Δ*p* around 2-10*kPa*. A deviation from these ranges results in structures that are either too thick or too thin to exist in intermediate stages of cell plate development based on literature. An application on a single oblate spheroid structure is shown in Fig. 3, and a summary of the full range of parameter values are given in Table 1.

**Table 1.** Model Parameter Ranges

| PARAMETER |  | VALUE |
| --- | --- | --- |
| Bending Modulus | $K_B$ | 62.5-200pNnm |
| Spontaneous Curvature | $c_0$ | 0-0.04nm<sup>-1</sup> |
| Pressure Difference | $\Delta p$ | 2-10kPa |
| Spreading Force Parameter | $\lambda$ | 0-6pN/nm |
| Gaussian Bending Modulus | $K_G$ | -0.8K<sub>B</sub> |
| Surface Tension Parameter | $\gamma$ | 1.6pN/nm |

**Figure 3.**
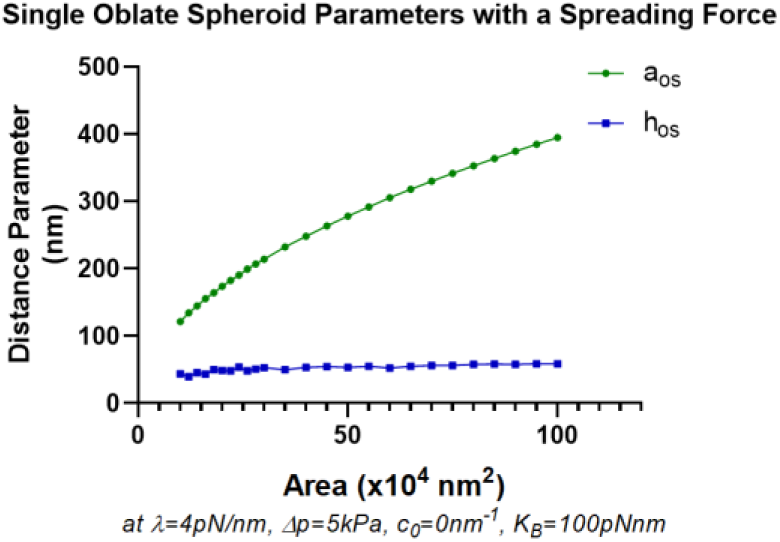
Evolution of single oblate spheroid parameters in the presence of a spreading force. Here, *h*_*os*_ = 2*c* (*a*_*os*_, *c* shown in Fig. 2*A*), thereby representing the overall height, or thickness, of the oblate spheroid. In the presence of a spreading force, we see that the thickness of the oblate spheroid remains in the 40-80nm range despite the increase in area. This reflects the thicknesses and growth patterns found in large vesicles in intermediate cell plate stages (1). In the absence of a spreading force, *h*_*os*_, or the thickness, would grow to values that are not observed in experiment. For reference, an area of 10^4^*nm*^2^ is roughly equal to that of a single vesicle.

Although the range of values for the pressure difference and the planar spreading force parameter were phenomenologically determined, they are within reasonable bounds. For instance, the solute concentration difference between the interior and the exterior of the cell plate by employing the van’t Hoff equation, which yields a solute concentration difference between 8×10^−4^*mol*/*l* and 4×10^−3^*mol*/*l*, comparable to protein solute concentration differences in higher plant cells. The spreading force due to callose over a length of a nanometer is around 2-6pN, which is comparable to the polymerization ratchet forces of a microtubule(25).

The latter force of course derives from stochastic addition of polymer at the growing tip of a polymer, which is not the physical situation here. We presume that the spreading force derives from the growth of callose polymers produced at individual callose synthase sites in the membrane drawing from the cytosolic supply of glucose. As outlined in the supplemental material, this does provide a basis for understanding the origin of the coefficient *λ* for the spreading force, in analogous to derivations of pressure from elementary kinetics. Namely, if we assume that the manufactured polymer has mean square end-to-end excursion described by the Flory self-avoiding polymer theory in two-dimensions (26) 

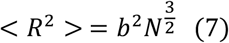

where *b* is the polymer link length, of order the persistence length for the polymer, and *N* is the number of polymers, then it follows that the spreading force coefficient identified as the work per unit area exerted on the boundary of the polymer disk is given by 

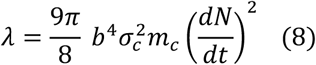

where *σ*_*c*_ is the density per unit area of the polymer, with monomer mass of *m*_*c*_. Using *b*∼10 *nm*, and a reasonable value for *σ*_*c*_, this provides an estimate of 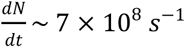 for 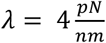. This of course represents the production of all the polymer synthases in the cell plate, and it is unknown what the production rate per synthase is. Literature values for individual cellulose synthase enzymes range from 1-10 cellulose links/s (see for example (27)), and values that low would require unphysically high numbers of synthesizing enzymes on the plate. However, we cannot easily connect any of these results to callose since essentially nothing is known about the morphology of callose within the plate or about the density or synthesis rate of callose synthase enzymes on the plate.

Our goal was to assess within the modeled free energy of Eq. (1), as to whether a spreading force is essential for the necessary transitions from a combination of tubulo-vesicular network to a fenestrated sheet and finally to a single mature cell plate structure. From an energy perspective, we identified that in the absence of a spreading force, single tubulo-vesicular and fenestrated structures have a lower value of energy at minima and are more stable than a single late stage cell plate structure (resembled by a single oblate spheroid) of the same area. Fig. 4*A* shows the energy minima of some tubular and fenestrated structures (7×6×0,…) compared to those of a single oblate spheroid (1×0×0). Δ*E*_*min*_ represents the difference of the energy minima value of the labelled structure with that of a single oblate spheroid, so that Δ*E*_*min*_(2×1×0)= *E*_*min*_(1×0×0)−*E*_*min*_(2×1×0). Thus, positive values indicate the relative stability of the labelled conformation. The general trend seems to indicate that in the absence of a spreading force, increasing tubularity is preferred with the increase in area. Interestingly, some fenestrated structures are more stable than a single oblate spheroid at the same area as well. Experimental results confirm that in the absence of callose, the maturation of the cell plate is arrested. Treatment with ES7, which inhibits callose deposition at the cell plate (12), leads to an abnormal transition of cell plate development such that it fails to mature into a cross wall. Fig. 5*E, F* show an example of arrested cell plate development in the absence of callose, in contrast to gradual cell plate expansion concomitant with callose deposition as in Fig. 5*A, B*. Cumulatively, the data supports the model in which callose provides the spreading force and mechanical support necessary for cell plate development.

**Figure 4.**
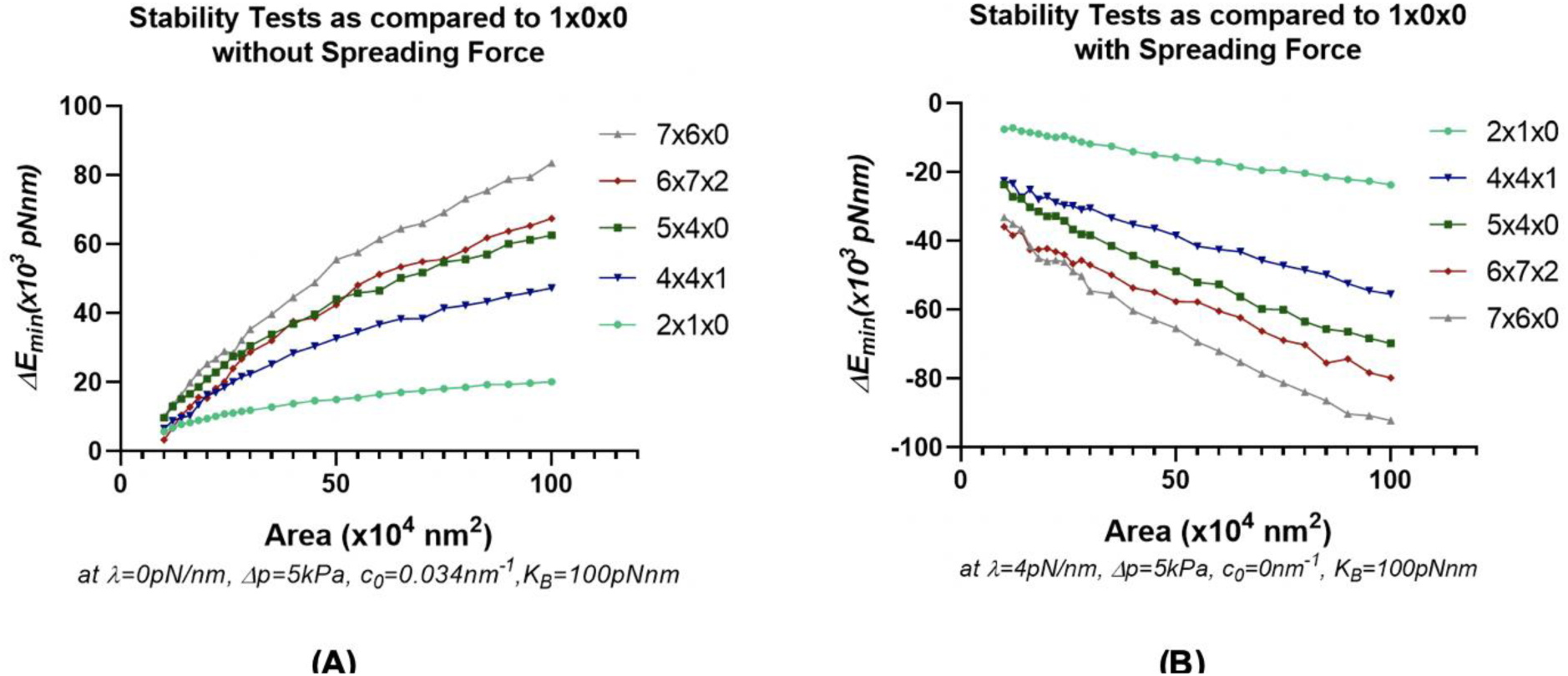
Stability tests determined by Δ*E*_*min*_ vs. Area for different conformations as compared to a single oblate spheroid at the labelled area. Here, a positive value of Δ*E*_*min*_ indicates relative stability of the labelled conformation as compared to a single oblate spheroid (1×0×0). (*A*) Relative stability of tubular and fenestrated structures in the absence of a spreading force with a finite spontaneous curvature. (*B*) Stability of a single oblate spheroid over tubular (2×1×0, 5×4×0, 7×6×0) and fenestrated (4×4×1, 6×7×2) structures in the presence of a spreading force and with zero spontaneous curvature. Note that in (*B*) a decrease of spontaneous curvature to a threshold value close to 0.015*nm*^−1^ yields similar results.

**Figure 5.**
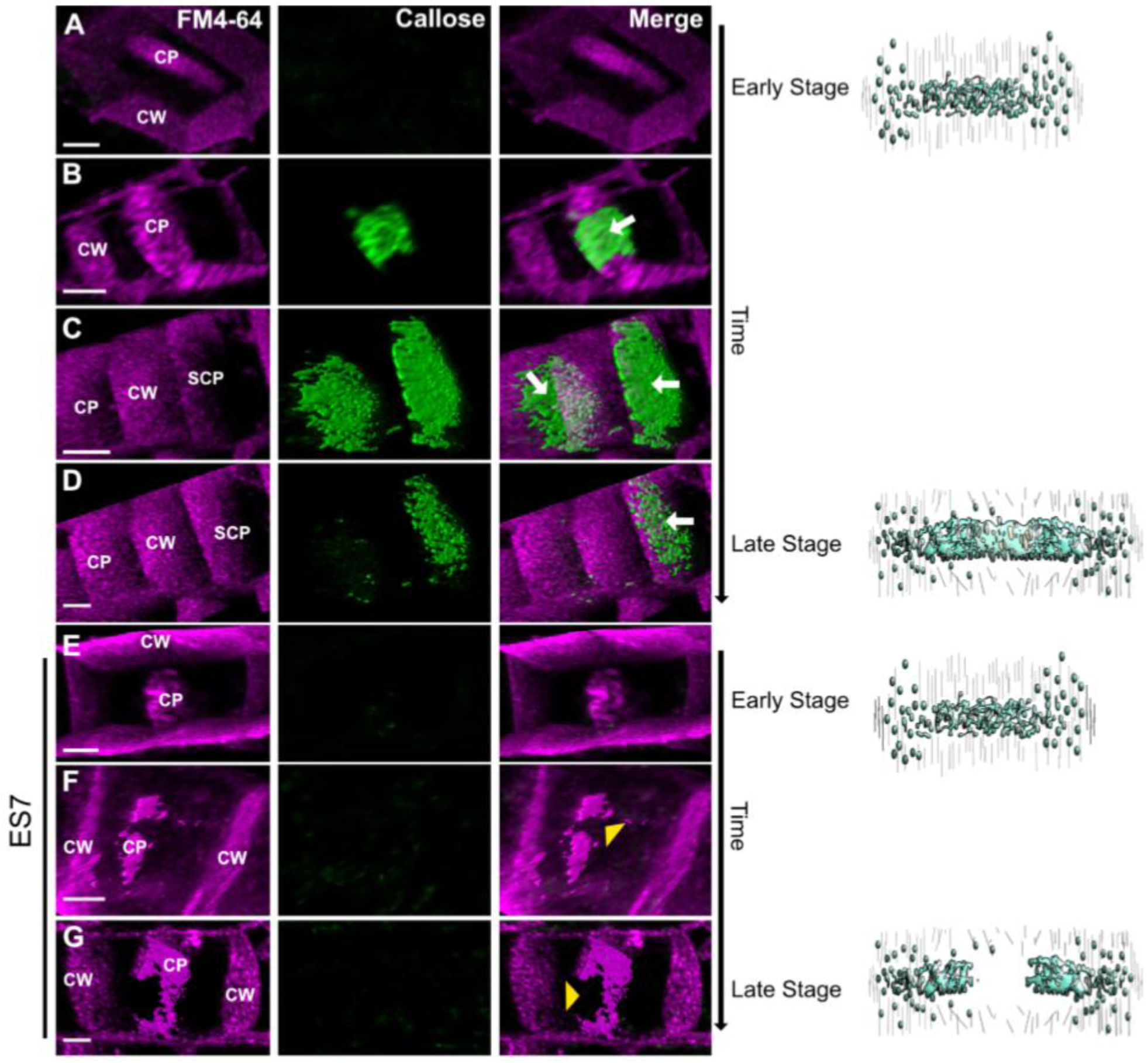
Progression of the cell plate in the presence and absence of callose. (*A-D*) Cell plate progression in the presence of callose. (*A*) shows an early stage cell plate before the accumulation of callose, similar to that depicted in Fig 1*A, B*, while *B-D* represent later cell plate stages including sheet like cell plates (SCP) as indicated in Fig 1*D*. Cell plate progression is shown by plasma membrane staining by FM4-64 (magenta), and callose accumulation staining with Aniline Blue fluorochrome (green). Note the transient accumulation of callose in later stages leading to the maturation of cell plate during normal cytokinesis (*B-D*). In *C*, two cell plates can be seen, and as maturation continues to *D*, callose is removed from one cell plate indicating its transition to a mature cell plate. (*E-G*) depict the progression of cytokinesis under ES7 that inhibits callose deposition. Note, early cell plate development is not affected with ES7 (*E*). However, in late stages of cell plate development under ES7 treatments, the absence of callose prevents the transition of membrane network into a stable single structure, leading to characteristic ‘cell plate stubs’ (*F* and *G*). CP, indicates cell plate, SCP, indicates sheet-list cell plate as depicted in Fig. 1*D*. Arrows indicate callose accumulation at the cell plate. Yellow arrowheads denote lack of callose at cell plate breakage points. Scale bar = 3µm

In the presence of a spreading force, we examined the possibility of a transition from tubulo-vesicular networks to a single oblate spheroid. Within the theory and the variational approach, we find that this is possible if the spontaneous curvature decreases to a threshold value (about 0.015*nm*^−1^) with larger cell plate area, i.e. in the presence of a spreading force. From an energy perspective, that would mean that tubulo-vesicular networks, as well as fenestrated sheets, should be unstable as compared to a single oblate spheroid, thereby preferring to change their morphology to one that is without tubes or fenestrations. While a finite spontaneous curvature is necessary to explain the origin of stability of the incoming vesicles (28), one can expect a change in the spontaneous curvature of the membrane due to the expected changes in membrane composition and protein activity that occurs during cell plate development (29). Fig. 4*B* shows the relative instability of selected tubular and fenestrated structures as compared to a single oblate spheroid in the presence of a spreading force and zero spontaneous curvature. Less tubular structures are now energetically favorable than highly tubular or fenestrated structures, with a single, complete structure being the most favorable. This represents the transition of a vesicle network to an expanding and mature cell plate (Fig. 5 *A* and *B-D*) in the presence of callose. It is noteworthy that callose accumulation starts at the center of the cell plate (Fig. 5 *A, B*, arrow), overlapping with later cell plate stages that transition from a vesicular network to a fenestrated sheet. For structures with genus zero (such as 2×1×0 or 3×2×0), we can also map a path to a single oblate spheroid if we relax the parameter restrictions that were initially imposed during the variational calculation. Fig. 4*B* shows results with the parameter restrictions in place. Fig. S3 shows data for a fenestrated structure in the absence (Fig. S3*A*) and the presence (Fig. S3*B*) of a spreading force, leading to fenestration shrinkage with the parameter restrictions in place.

To better represent a biological system, we compared multiple 2×1×0 structures (approximating accumulated fused vesicles forming a network) to a single oblate spheroid of the same combined area. Similar to our earlier calculations, we find that in the absence of a spreading force, a single oblate spheroid is less stable, as in Fig. 6*A*. The relative instability is magnified with the increase of area, and with the increase in the number of tubular structures. In the presence of a spreading force and a decreased spontaneous curvature, as in Fig 6*B*, the inverse is true, favoring fewer complex structures. When comparing multiple structures to a single mature structure of the same area, there is no need to enforce the decrease in spontaneous curvature. However, for consistency, results with a zero spontaneous curvature in the presence of a spreading force, and a finite spontaneous curvature in the absence of a spreading force are shown.

**Figure 6.**
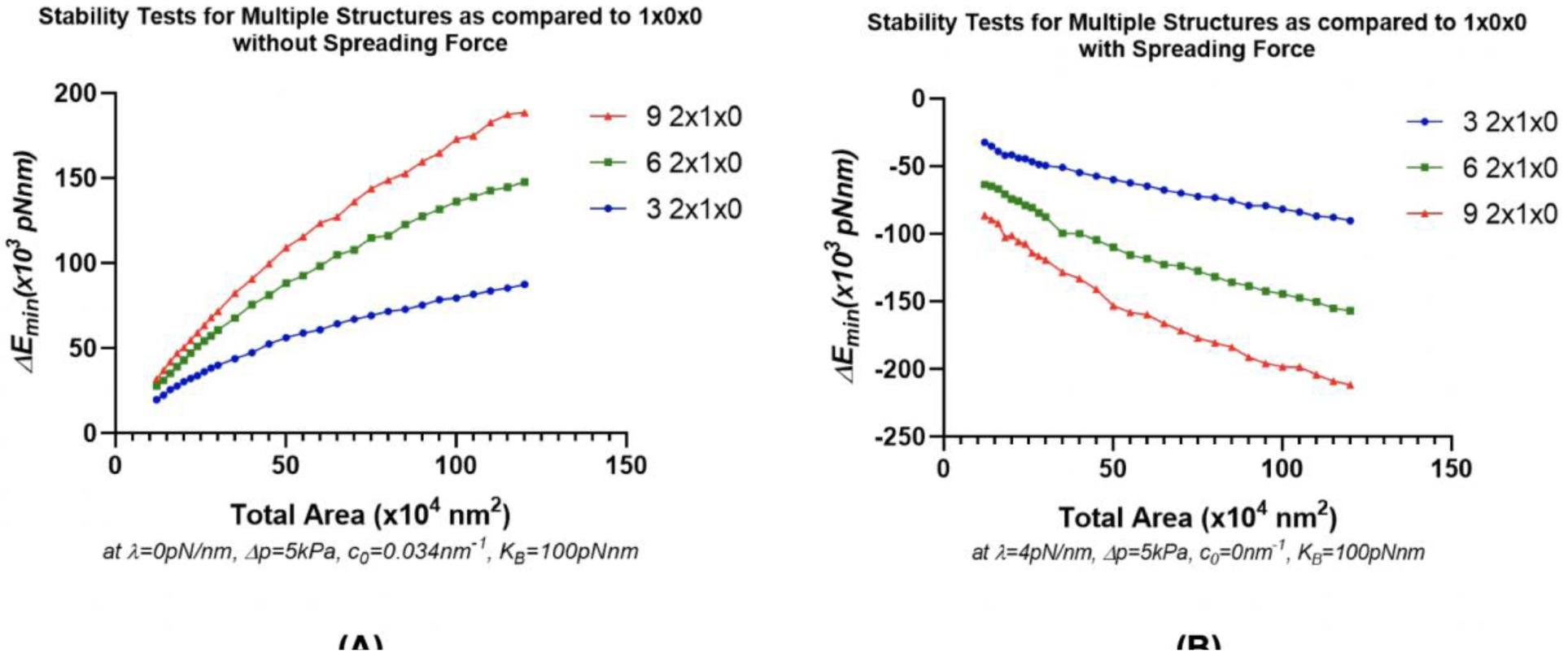
Stability test for multiple 2×1×0 structures as compared to a single oblate spheroid at the labelled area. (*A*) Relative stability of multiple 2×1×0 structures as compared to a single oblate spheroid in the absence of a spreading force. At a labelled area, a larger number of structures have a higher, more positive value of Δ*E*_*min*_ collectively, thereby indicating that in the absence of a spreading force, tubular structures as well as emerging fenestrated/network structures (as inferred by the results of Fig 4.) would tend to accumulate as in Fig 1*C*. (*B*) Stability of a single oblate spheroid as compared to multiple 2×1×0 structures in the presence of a spreading force and with zero spontaneous curvature. At a labelled area, a larger number of structures have a lower, more negative value of Δ*E*_*min*_ collectively, thereby indicating the energetic favorability of structures fusing to form larger, more mature structure(s) in the presence of a spreading force.

The exact values of pressure difference, spreading force, and spontaneous curvature that are needed for a transition to a complete cell plate structure depend on the choice of bending modulus. Additional calculations with different choices of bending modulus are given in the supplemental figures (Figs. S4-S8). In general, a stiffer membrane, one that is represented by a larger bending modulus, requires a stronger spreading force in addition to a higher pressure difference to transition to a mature cell plate structure. By testing over the full range of literature values for the bending modulus, we also account for the possible differences in the bending modulus that may arise in different regions of the cell plate due to varying thickness and rigidity. However, as shown in Figs. S4-S8, the need for a spreading force is always there, regardless of the choice of the bending modulus. We also show calculations for larger, highly tubulated fenestrated structures in Figs. S9-S10, which are stable in the absence of a spreading force, but unstable as compared to a single oblate spheroid in the presence of a spreading force, further supporting our hypothesis.

## Discussion

While other force generating proteins, including but not limited to dynamin-like springs and clathrin, are involved in the tubulation and membrane material recycling processes(30-32), the hypothesized planar spreading force ensures that the resulting tubulo-vesicular and fenestrated structures do not collapse onto themselves, thereby enabling them to transition to more mature cell plate structures. This is evidenced in Fig 4*A*, where we show that existing fenestrated and tubulo-vesicular structures are more stable compared to a single mature cell plate structure of the same area in the absence of a spreading force.

From an energy minimization analysis, we have shown that a planar spreading force is vital for cell plates to transition from tubular and fenestrated networks to complete, late stage/mature cell plate structures. We also show that in the absence of a spreading force, the addition of membrane material yields stable tubulo-vesicular and fenestrated structures, but that those structures are unable to mature beyond that. Given that spatio-temporal experimental data in the absence of callose shows the failure of cell plates to mature beyond tubular and fenestrated networks, our model supports the hypothesis that callose provides a spreading force facilitating these transitions. The need for this spreading force is magnified when we compare single cell plate structures to multiple smaller structures of the same total area.

We still do not know what the precise molecular scale mechanism for this spreading force is. The development and adaptation of advanced analytical methods to study biomechanics at subcellular resolution can shed light on the nature of this spreading force and its regulation.

As noted in the results section and the supplemental information, expansion of a two-dimensional self-avoiding polymer gives a crude areal pressure expression of the correct form and a connection to the rate of polymer synthesis. This two-dimensionality would arise from the internal tethering of the polymer chains to the inner membrane surface of the plate, thereby resulting in a effective planar spreading force. The presence of callose near the surface rather the center of the forming cell plate (3), especially in later cell plate development stages, further supports the hypothesis that the primary relevant polymer is callose. This raises the intriguing possibility of a common origin to the decrease in spontaneous curvature and onset of a spreading force. In the model results, the spreading force is relevant when there is sufficient connection of individual oblate spheroidal vesicles, and it is at this stage that we shut off the spontaneous curvature. The nanoscale surface topography can potentially serve as a direct biochemical signal to activate this process (33). The possible tethering of the callose to the membrane could concomitantly induce spreading and reduce spontaneous curvature by modifying the membrane mechanics.

Callose is produced in β-1-3 linear glucan polymers that show a low degree of crystallinity(34). The proposed conformation of inter-molecular associated forms of callose that include triple helices along with loop regions supports a semi flexible network formation(35, 36). The properties exhibited by callose aggregates in such a network are ideal for generating structural molds that can effectively produce planar forces. Detailed experiments are necessary to determine how the possible callose conformations contribute to different magnitudes of spreading force, an analysis which is the focus of our future studies. Callose could also serve as a scaffold into which other more permanent polysaccharides and proteins are later deposited(34, 37). Callose aggregates would allow insertion or co-gelation with other polymers at the cell plate which together may contribute to the spreading force and change of curvature. The properties of callose as a plasticizer allowing for deformation without breakage as opposed to pure cellulose(38) support its transient accumulation during the transition of a membrane network stage to a fenestrated sheet and a mature cell plate. Furthermore, the potential transient interaction with cellulose may also contribute to a flexible yet strong polymer that supports the maturation of the cell plate(38). Taken together, our model provides a basis for understanding how membrane structures evolve in the presence of a spreading force and will likely shed light in such transitions that occur beyond cytokinesis.

## Author Contributions

D.C. and G.D. designed research. M.Z.J. did computational work and R.S. did experimental work. M.Z.J. wrote the paper, with significant revisions from D.C. and G.D. R.S. contributed to the materials and methods section, as well as figure illustrations and captions.

## Acknowledgments

This work was supported by NSF Grant MCB 1818219. We would also like to thank the members of the Drakakaki lab, particularly Destiny Davis and Michel Ruiz for useful discussions pertaining to the application of the model and our understanding of the problem. The authors declare no competing financial interest.

## Supplementary Information

### Supplemental Material

#### Parameter Set Up

To implement energy minimization on a parameterized basis set defined at a particular surface area, we first identified the parameter space for a given conformation that yielded the desired area. For a single oblate spheroid, only two parameters are needed, namely, the radius along the major axis, ‘*a*’, and the radius along the minor axis, ‘*c*’. However, for the consolidated tubular networks and emerging fenestrated structures, one must account for continuity between the oblate spheroid and the elliptic hyperboloid, while also making the relevant corrections in area. This continuity can be achieved by matching the slopes of the elliptical hyperboloid with the oblate spheroid along the primary axes, while enforcing contact.

A normal one-sheeted elliptical hyperboloid centered at a distance *d* along the x axis can be described by the following equation: 

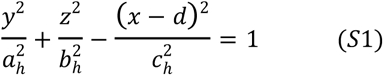

Here, *a*_*h*_ is the skirt radius along the xy-plane, *b*_*h*_ is the skirt radius on the xz-plane, and *c*_*h*_ describes the elongation along the y-axis, where the xy-plane defines the equatorial plane, and the z-axis is the polar axis. Fig. S2 shows these parameters for a hyperboloid in a 2×1×0 structure.

By enforcing continuity with an oblate spheroid at the origin, we can derive the following relationships for *d* and *b*_*h*_ such that: 

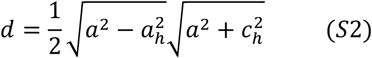

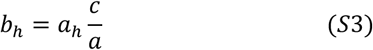

Using this, the length of the hyperboloid *l* is given by: 

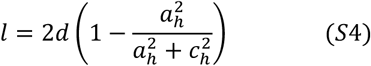

Using these restrictions, we can form connected structures of oblate spheroids and hyperboloids to form approximate representations of tubular and fenestrated structures. Additional demands that are placed by specific conformations such as in complicated emerging fenestrated structures (such as 6×9×3) were taken into account by eliminating choices of hyperboloids that would cause clashes.

Thus, our complete parameter space for a given conformation at a particular area is given by a list of values of (*a, c, a*_*h*_, *l*), where the corresponding area is calculated by numerical integration methods up to an error tolerance of 0.01%, taking into account spatial constraints.

#### Calculation of area elements

To calculate the mean curvature and the area elements of the parameterized structures, we employed the following methods: 

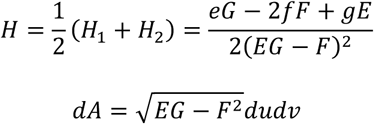

Where *E, F*, and *G* are coefficients of the first fundamental form (line element) and *e. f*, and *g* are coefficients of the second fundamental form (shape tensor), and *u* and *v* parameterize the surface. The selection of E,F, G are based standard practices as described in(1). Thus, we can calculate the *E*_*bending*_ from Eq. (1) for any conformation given the conformation type and its corresponding parameter space.

#### Results with full range of bending moduli

The supplemental data in Figs. S4-S8 show additional calculations in a range of bending moduli. A bending modulus of 62.5pNnm (about 15K_b_T) corresponds to the lower range of bending moduli corresponding to published data(2), while a bending modulus of 200 pN-nm (about 50K_b_T) corresponds to the higher range. It is important to note that the key outcome remains the same, a finite spreading force coupled with a decrease in spontaneous curvature is essential for a transition to a single, complete cell plate structure, regardless of the choice of bending modulus.

#### Fenestrated structures data at higher areas

The supplemental data in Figs. S9-S10 show Δ*E*_*min*_ calculations for different types of emerging fenestrated structure conformations at larger cell plate areas. In the absence of a spreading force, and with finite spontaneous curvature, larger, more tubulated fenestrated structures are more stable than a single cell plate. In the presence of a spreading force and with decreased spontaneous curvature, a transition to a single, mature cell plate structure is energetically favorable.

#### Emergence of Spreading Force from Two-Dimensional Self-Avoiding Polymer Physics

As a finite thickness polymer, the mean square extent of a polysaccharide like callose is subject to the law of self-avoiding polymers, *viz.* 

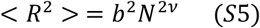

with, per Flory theory(3), 

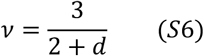

obtained from a balancing of two entropic effects of self-avoidance and entropic springiness (high probability of zero end-to-end distance). Here *b* is the size of a polymer link, of order the persistence length. In two dimensions, 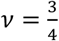. Hence, as the synthesizing enzyme produces more polymer, to the extent the polymer tethers to each side of the inner cell plate, we see that at the edge there will be a radial pressure due to the growing polymer front (Fig. S11). The radial growth speed *v*_*F*_ is given by 

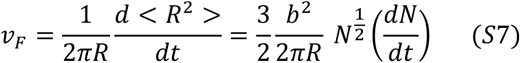

and the rate of change of area is 

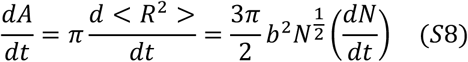

Hence, the magnitude of the radial force acting on the edge of the plate is (*m*_*c*_ is the mass of a constituent monomer, i.e., glucose, and *σ*_*c*_ is the areal density of polymer in the plate) 

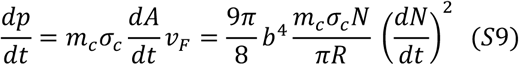

The spreading force *λ* or areal pressure is the total work done per unit area in expanding the plate a radial distance *dR* so 

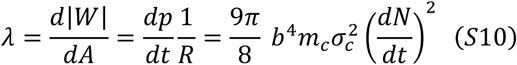

Hence, in this simple model, the spreading force is directly related to the production rate of polymer in the plate.

## Supplemental Figures

**Figure S1:**
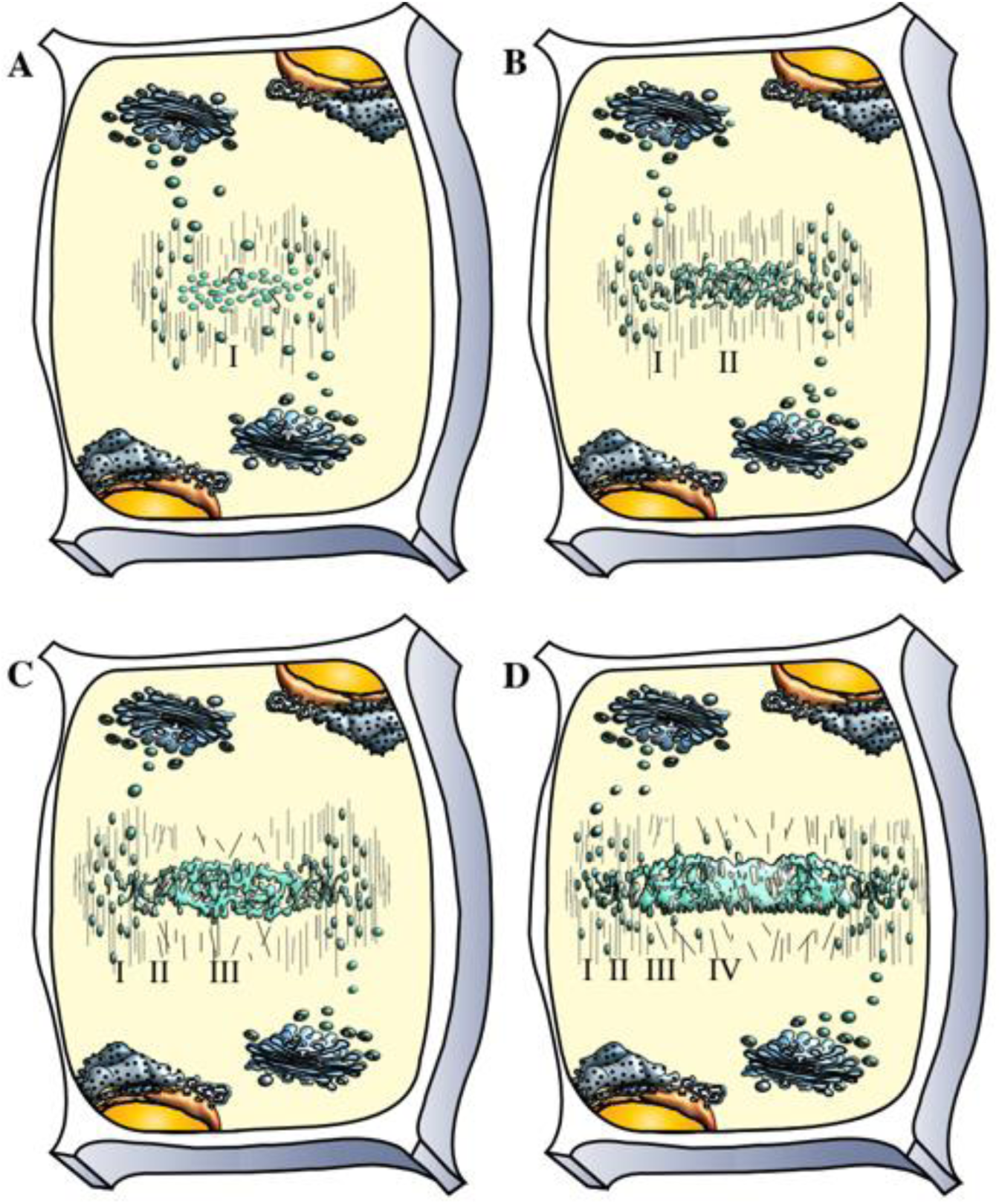
Schematic representation of cell plate assembly stages. Cell plate assembly occurs centrifugally in multiple stages. (*A*) During the first stage (I), cytokinetic vesicles guided by the phragmoplast accumulate at the center of the dividing cells, at the cell plate assembly matrix, where fusion starts to occur. (*B*) Vesicles undergo fusion and fission and conformational changes resulting in tubular-vesicular network (TVN) (Stage II). (*C*) Interconnected membrane structures transition to a tubular network (TN). At this stage high callose deposition occurs (Stage III). The membrane network further expands to an almost continuous fenestrated membrane sheet (PFS) (Stage IV), (*D*). Deposition of additional polysaccharides helps transition to a new cell wall, separating the two daughter cells. Note that different stages can occur simultaneously, images are not to scale. This simplified representation emphasizes on cell plate membranes(4, 5).

**Figure S2:**
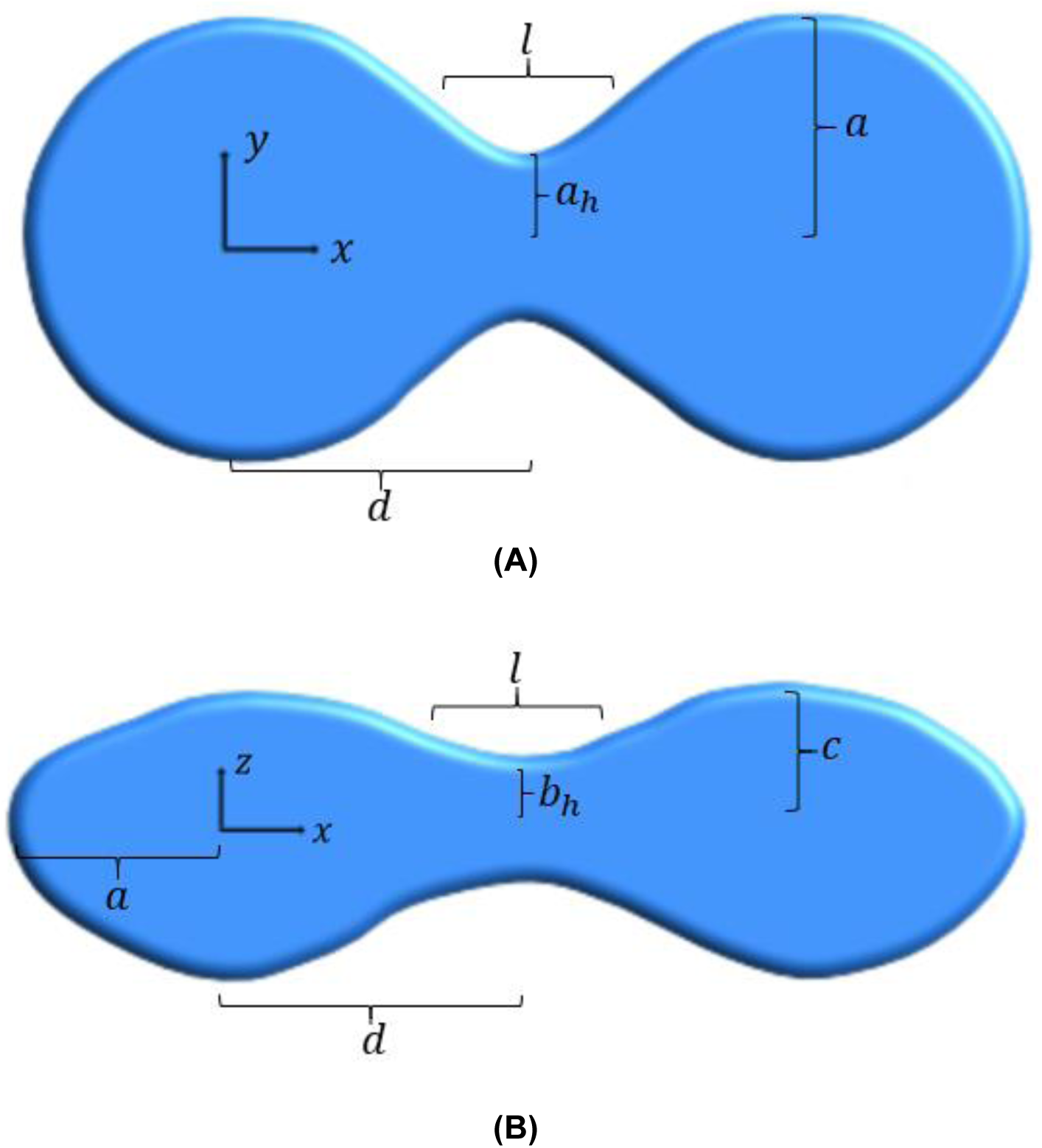
Parameters visualized on a representative 2×1×0 structure. To enforce continuity between an oblate spheroid of given parameters (*a, c*) and an elliptic hyperboloid with parameters (*a*_*h*_, *b*_*h*_, *c*_*h*_), we can calculate *d* and *b*_*h*_ are dependent variables, a full conformation can be described by the type of conformation and the parameter set (*a, c, a*_*h*_, *l*), or equivalently (*a, c, a*_*h*_, *c*_*h*_). The perpendicular arrows show the respective axes of the conformation. *(A)* shows the top view of the conformation, while *(B)* shows the side view of the same conformation.

**Figure S3.**
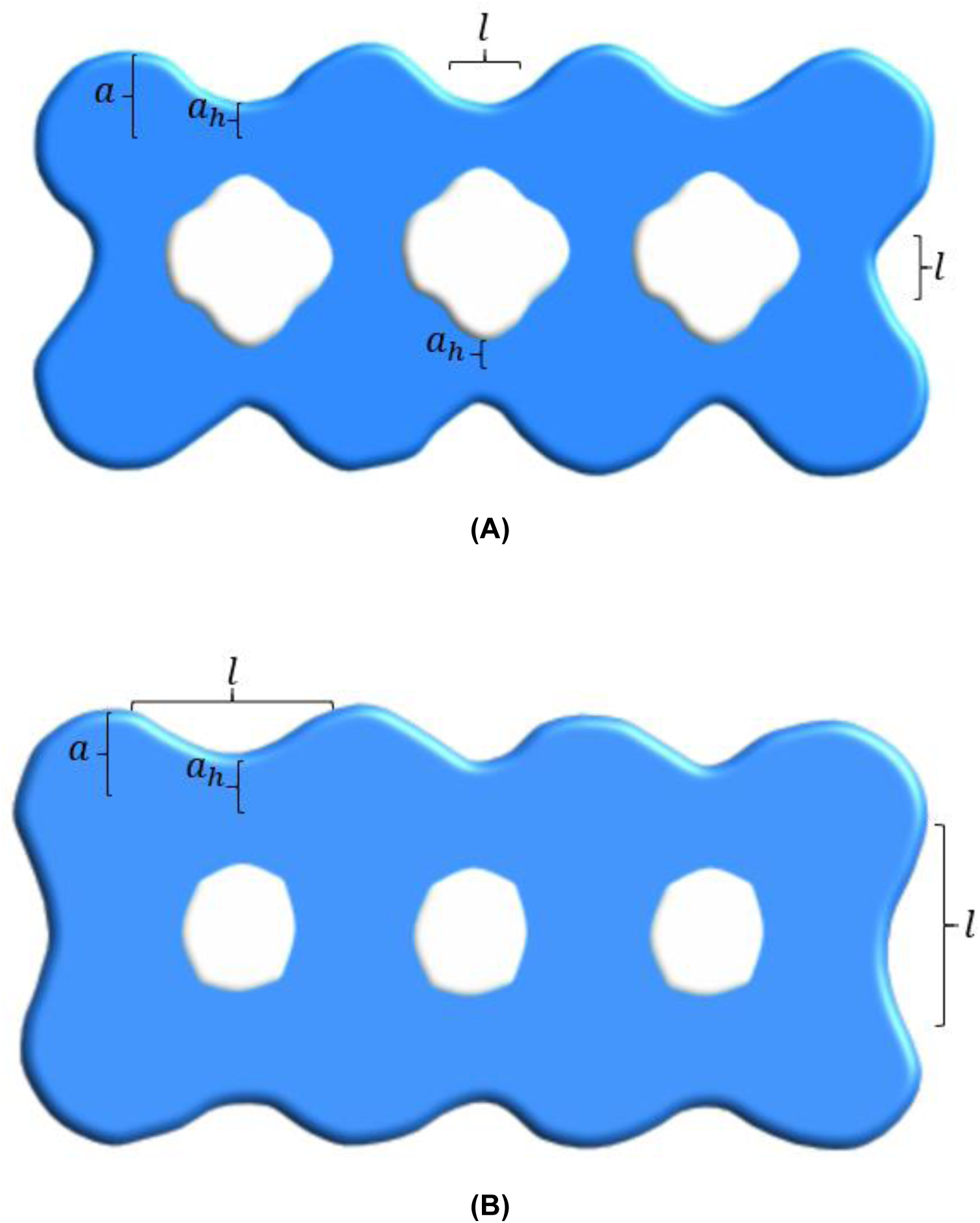
Effect of a spreading force visualized in a 8×10×3 conformation. The parameters shown here were extracted after energy minimization calculations on an 8×108×3 structure with parameter restrictions in place. In the absence of a spreading force, larger fenestrations, and narrower tubular connections are predicted, as shown in a top view in (*A*). This structure has an area of 2×10^5^*nm*^2^, while the parameters (*a, c, a*_*h*_, *l*) are given by (52,31.5,20,25.8)*nm*. As a spreading force is turned on and the spontaneous curvature is decreased, the tubular connections widen, thereby shrinking the fenestration sizes, as shown in a top view in (*B*). For the same area, the parameters change to (58,24,33.5,25.41)*nm*. If we relax the imposed parameter restrictions in the presence of a spreading force, the resulting structure would be of a single oblate spheroid with *a* = 173*nm, c* = 25*nm*.

**Figure S4.**
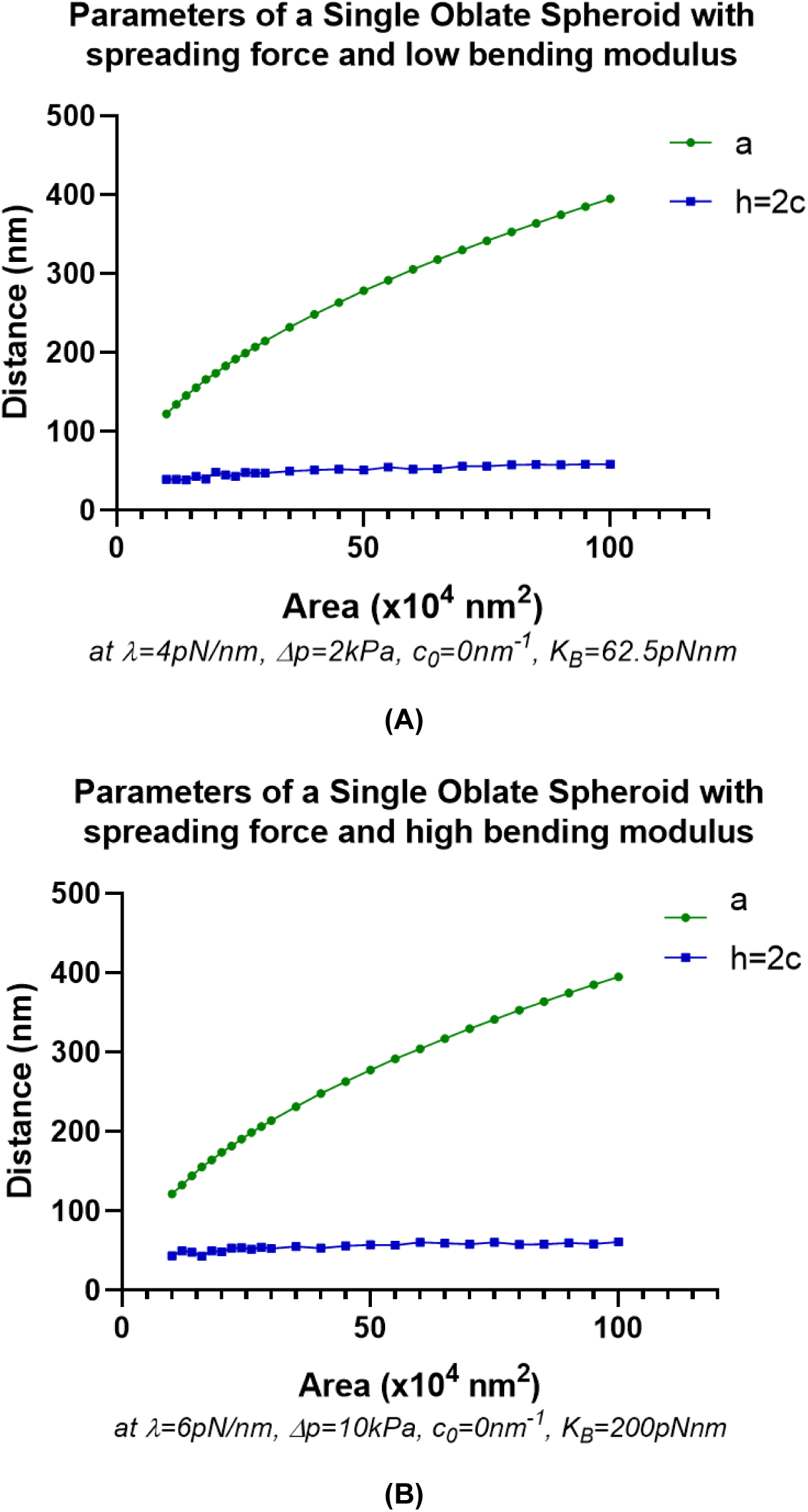
Evolution of single oblate spheroid parameters in the presence of a spreading force. Results with extremal values of the bending modulus are shown in *(A)* and *(B)*. Despite the increasing area, the height (h) remains in the 40-80nm region. With a smaller bending modulus, as in (*A*), a smaller value of the spreading force parameter *λ* and pressure difference Δ*p* is required to maintain the height within the desired region for the specified areas. With a larger bending modulus, as in (*B*), larger values of *λ* and Δ*p* are required.

**Figure S5.**
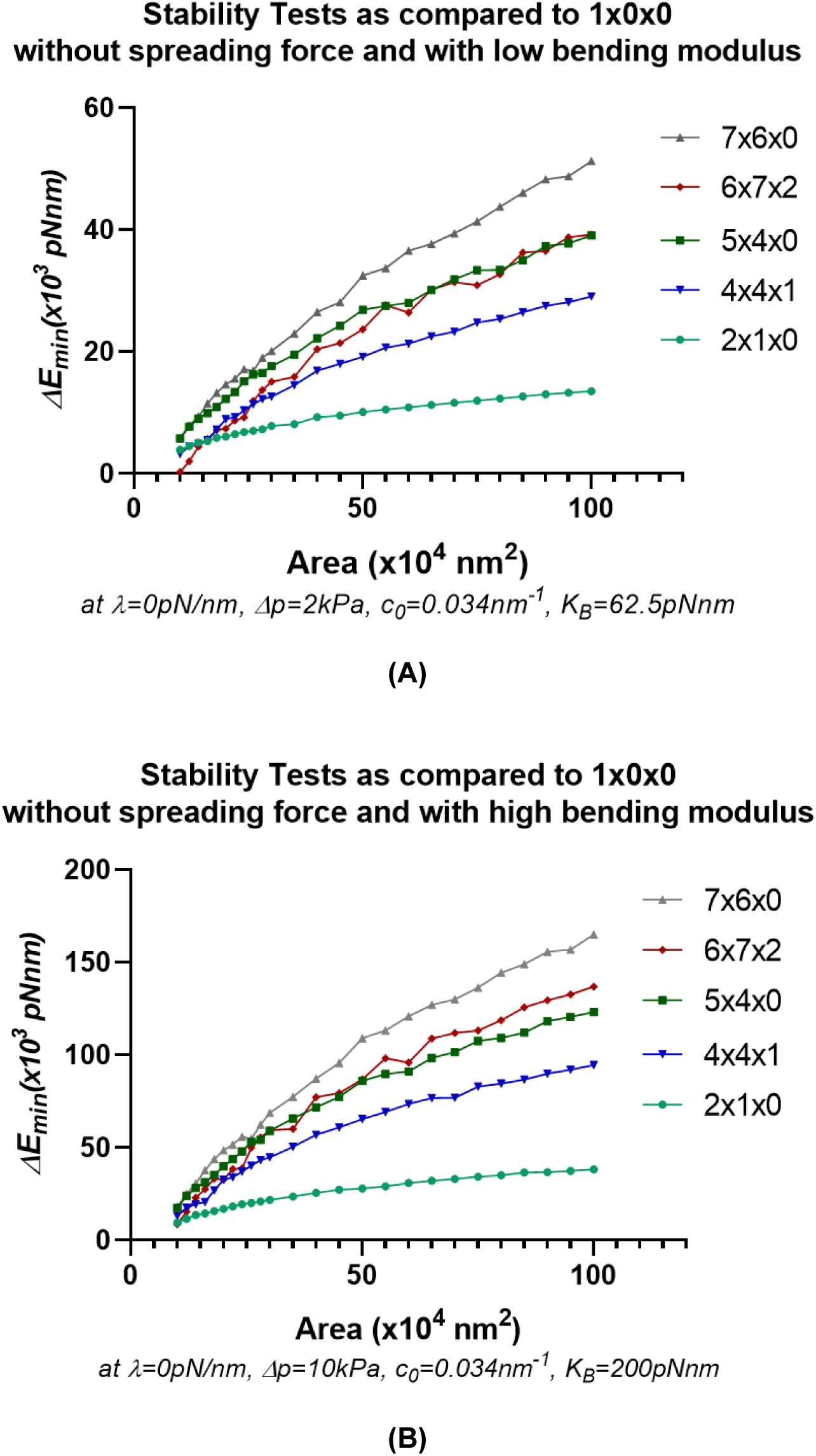
Stability tests of various configurations under different bending modulus in the absence of a spreading force. (*A*) Stability tests for a small value of bending modulus while (*B*) shows calculations for a larger value of bending modulus. A positive value of Δ*E*_*min*_ indicates relative stability of the labelled conformation as compared to a single oblate spheroid. Note that in the absence of a spreading force and finite spontaneous curvature, increasingly tubular and fenestrated structures are more stable as compared to a single oblate spheroid. The different values of Δp and *λ* arise due to the constraints on structure thickness as shown in Fig. 3 and Fig. S4.

**Figure S6.**
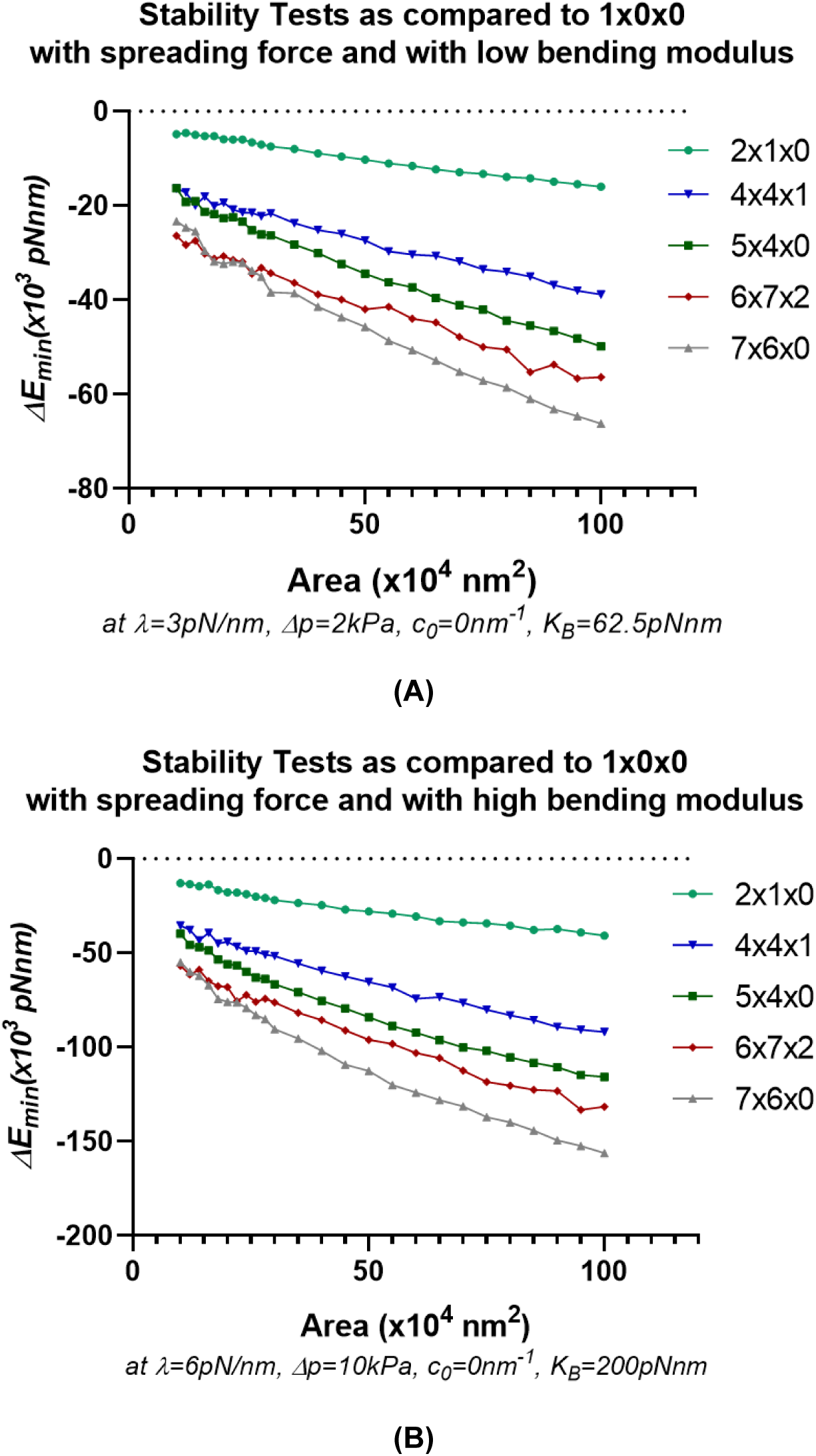
Stability tests of various configurations under different bending modulus in the presence of a spreading force and with zero spontaneous curvature. (*A*) Stability tests for a lower value of bending modulus and (*B*) higher value of bending modulus. Note that with the presence of a spreading force and with zero spontaneous curvature, increasingly tubular and fenestrated structures (i.e.7×6×0) are increasingly unstable as compared to a single oblate spheroid, indicating the energetic favorability for cell plate structures to mature to a disk like shape. A positive value of Δ*E*_*min*_ indicates relative stability of the labelled conformation as compared to a single oblate spheroid.The different values of Δ*p* and *λ* arise due to the constraints on structure thickness as shown in Fig. 3 and Fig. S4.

**Figure S7.**
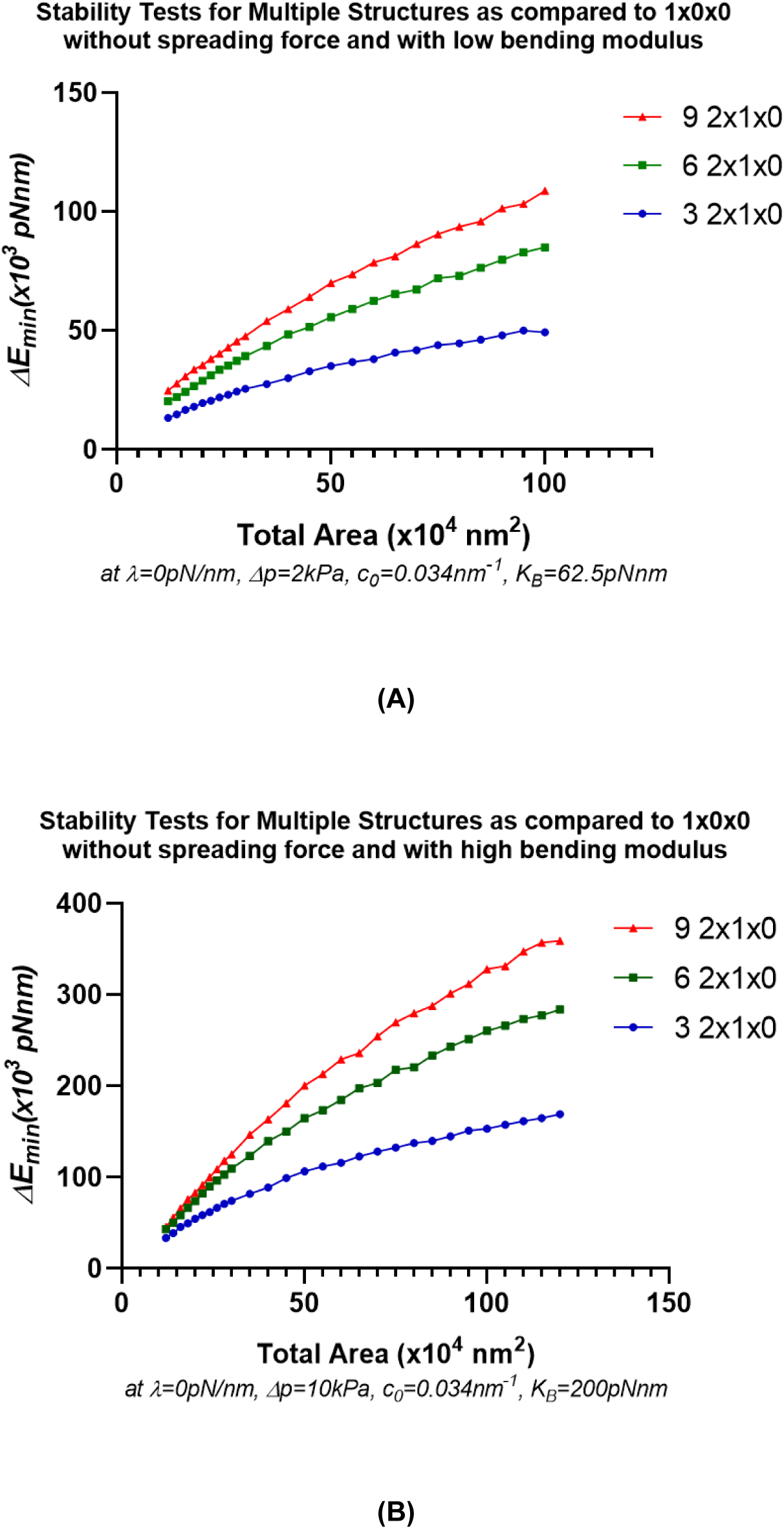
Stability tests of multiple 2×1×0 structures as compared to a single oblate spheroid in the absence of a spreading force and with finite spontaneous curvature. Δ*E*_*min*_ of multiple 2×1×0 structures as compared to a single oblate spheroid under for a low value of bending modulus (*A*) and high value of bending modulus (*B*) are shown. Note that in the absence of a spreading force and with finite spontaneous curvature, tubular structures are energetically favorable in these conditions, thereby modeling a membrane network stage. A positive value of Δ*E*_*min*_ indicates relative stability of the labelled conformation as compared to a single oblate spheroid. The different values of Δp and *λ* arise due to constrains on structure thickness as shown in Fig. 3 and Fig. S4.

**Figure S8.**
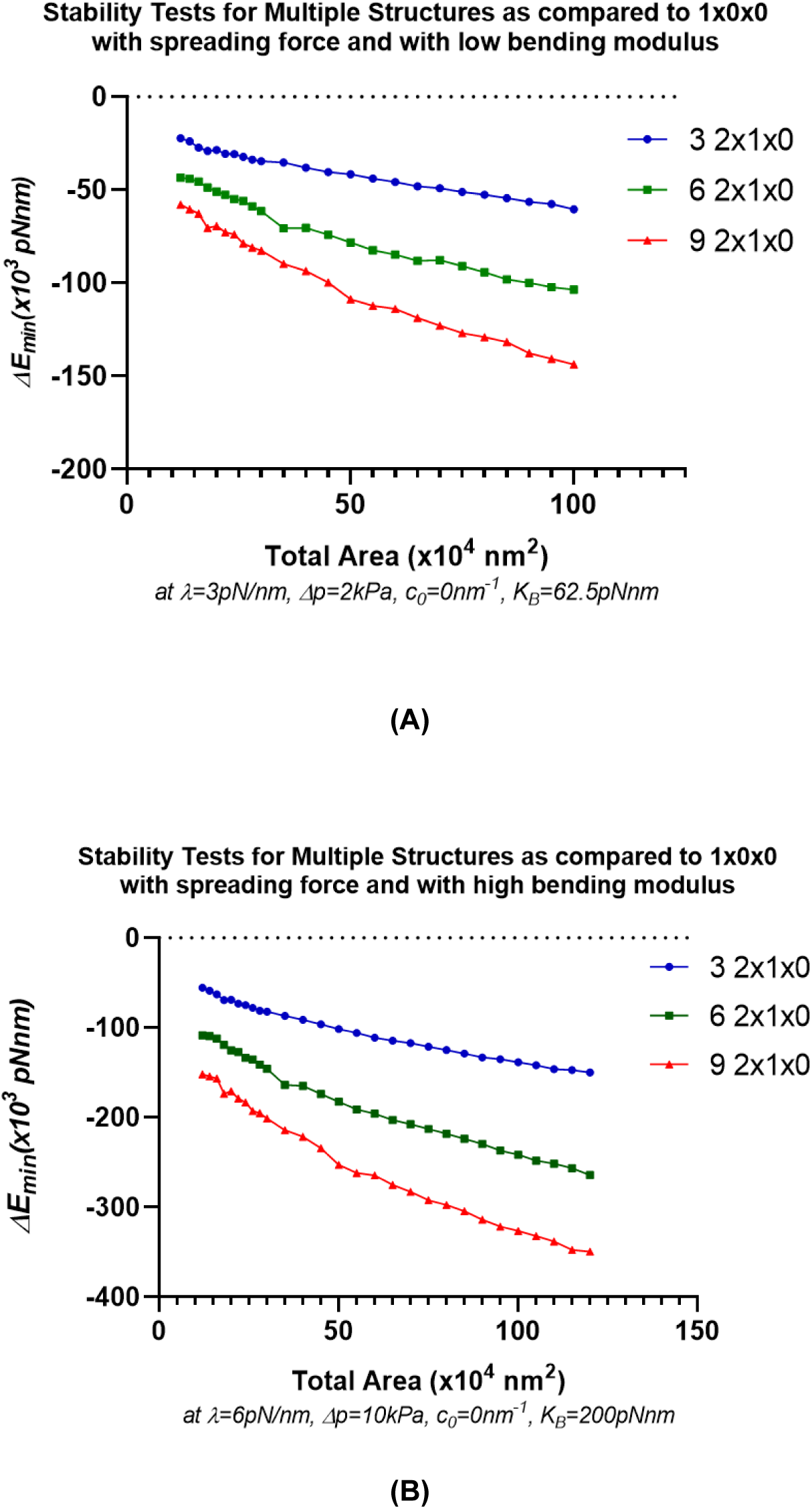
Stability tests of multiple 2×1×0 structures as compared to a single oblate spheroid in the presence of a spreading force and with zero spontaneous curvature. In the presence of a spreading force and with zero spontaneous curvature, tubular structures are unstable compared to a single oblate spheroid, thereby indicating the energetic favorability of structures fusing to form larger, more mature structure(s). (*A*) shows results for a small value of bending modulus while (*B*) shows results for a larger value of bending modulus. A positive value of Δ*E*_*min*_ indicates relative stability of the labelled conformation as compared to a single oblate spheroid. The different values of Δp and *λ* arise due to the limitations on structure thickness as shown in Fig. 3 and Fig. S4.

**Figure S9.**
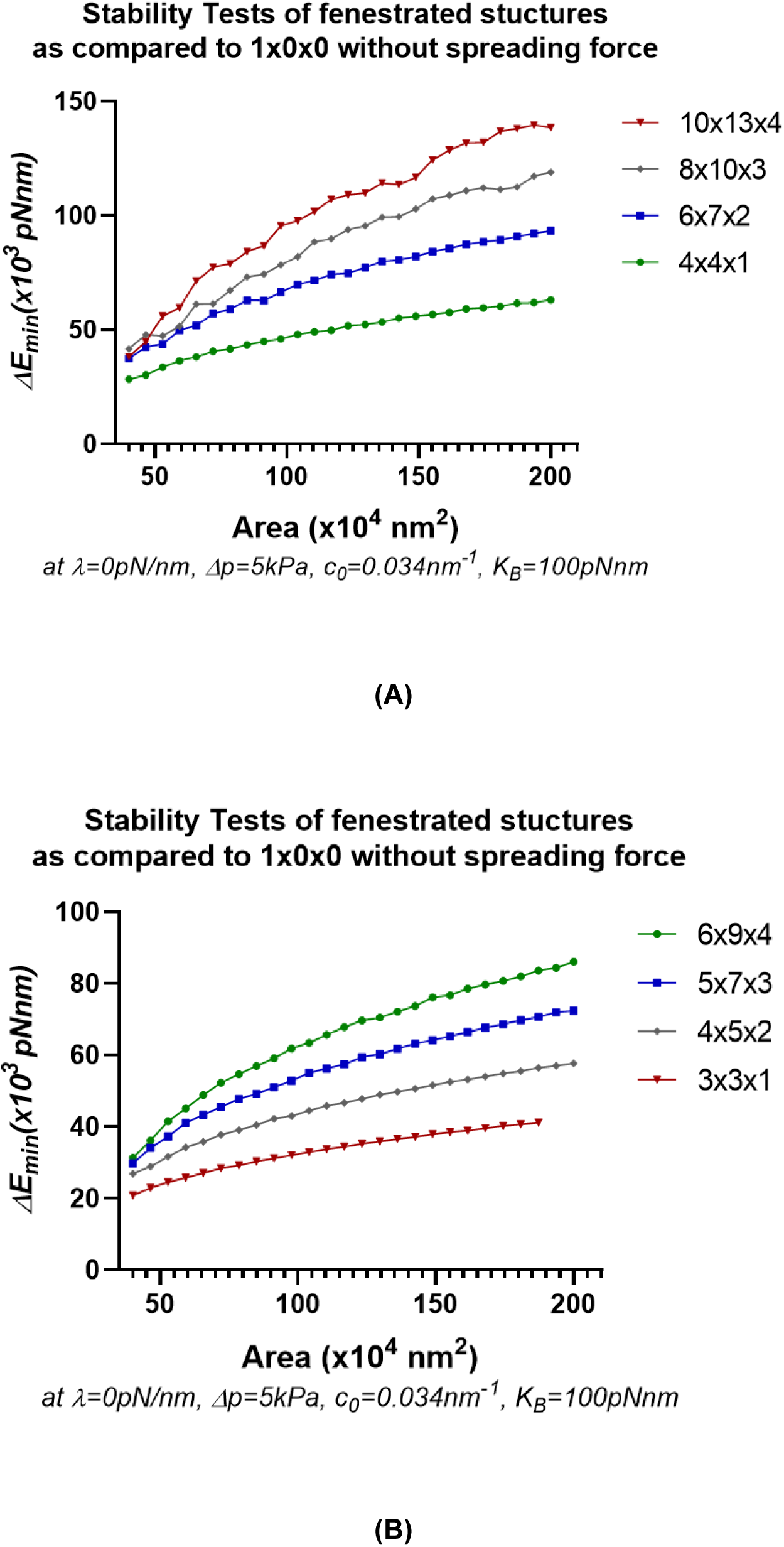
Stability tests of tubular/fenestrated structures as compared to a single oblate spheroid in the absence of a spreading force. In the absence of a spreading force, and with finite spontaneous curvature, fenestrated and tubular structures are, in general, more stable than a single oblate spheroid. This relative stability is magnified with the increase in area particularly for heavily tubular structures (10×13×4 in *A*, 6×9×4 in *B*), consistent with observations at tubular network/very early fenestrated sheet stages.

**Figure S10.**
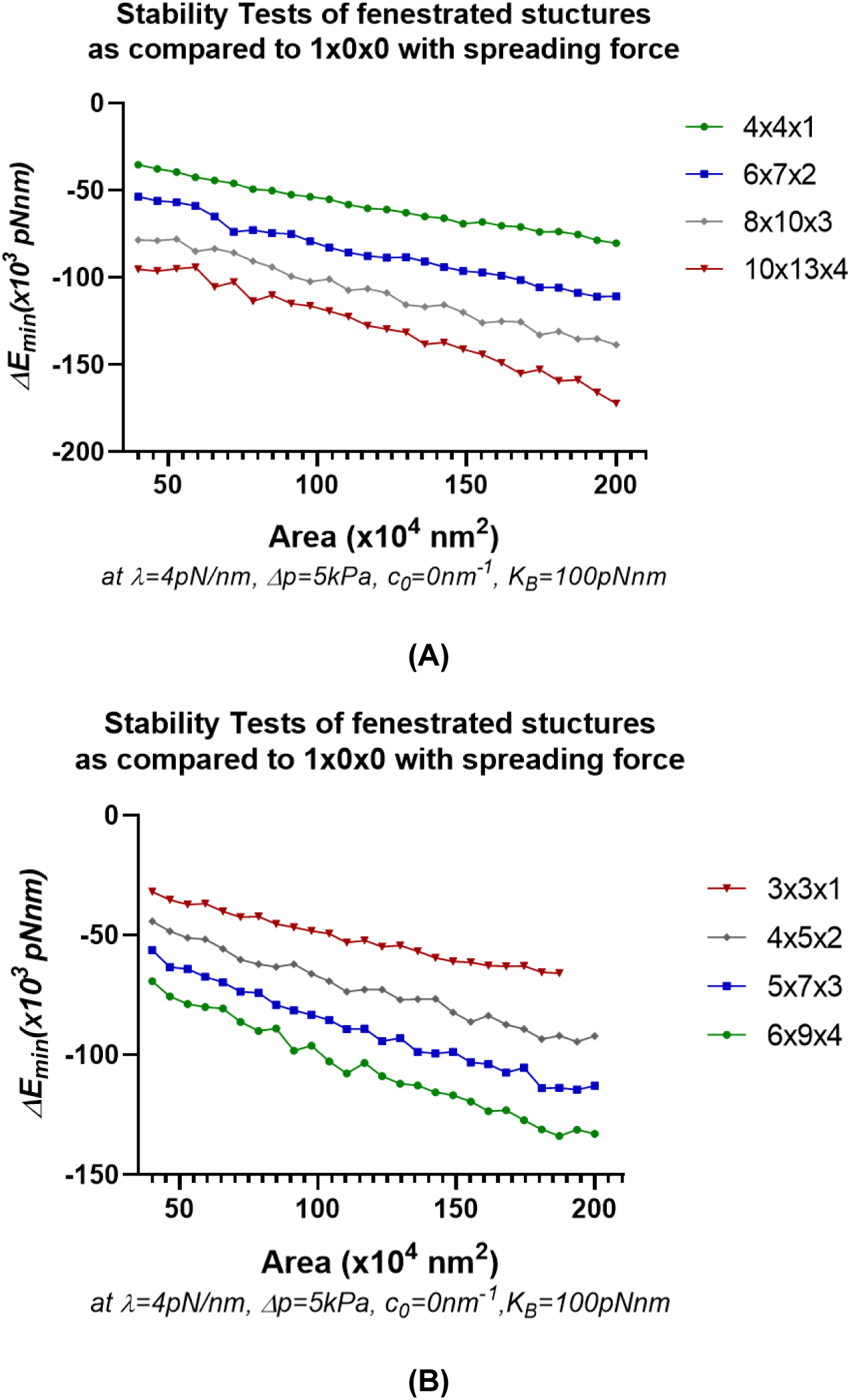
Stability tests of tubular/fenestrated structures as compared to a single oblate spheroid in the presence of a spreading force. (*A, B*) In the presence of a spreading force, and with decreased spontaneous curvature, a single oblate spheroid is more stable compared to larger, tubular, fenestrated structures. This indicates the necessity of a spreading force to incur a transition from a tubular/ fenestrated sheet stage to a single mature cell plate structure.

**Figure S11.**
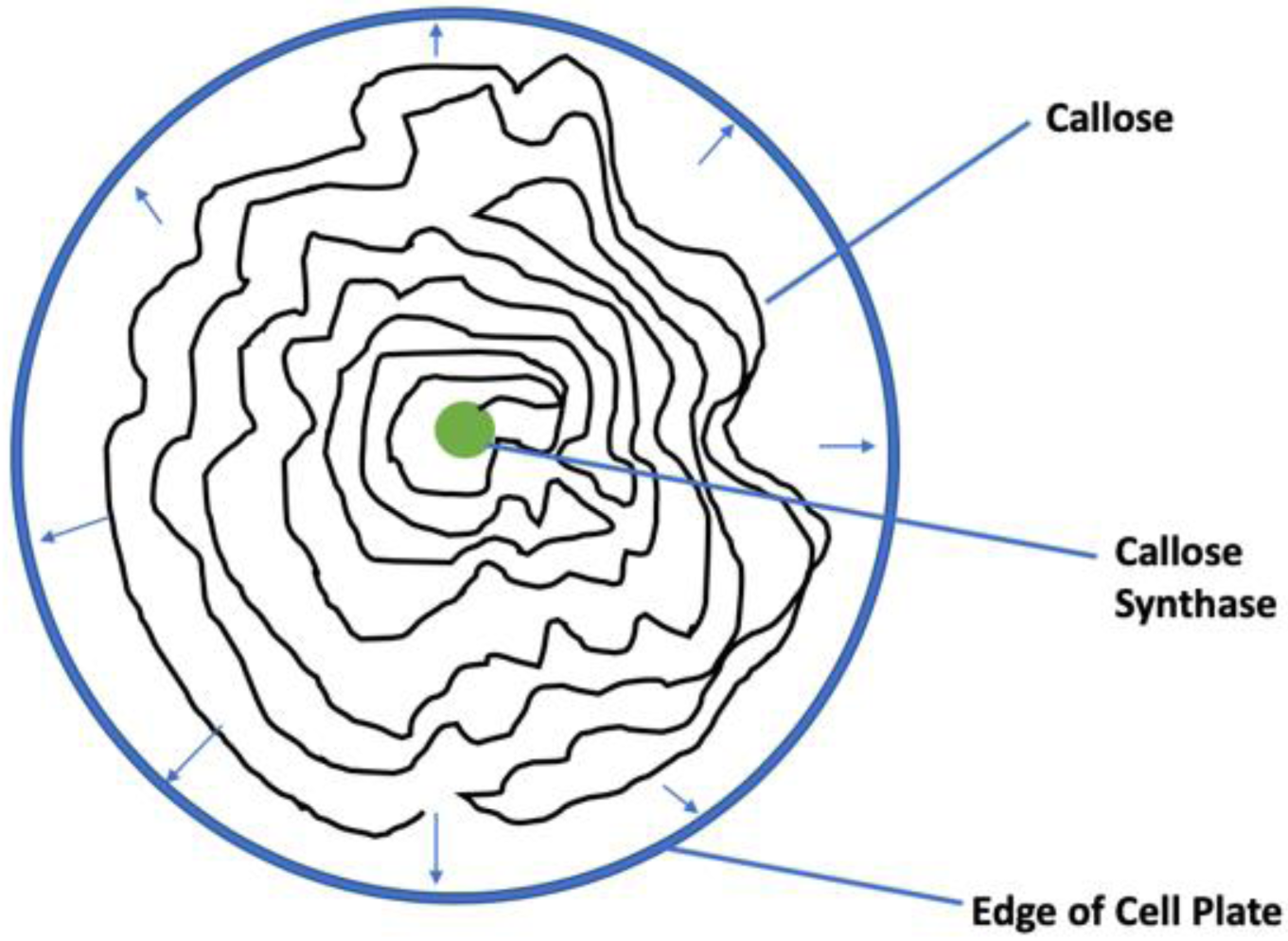
Growth of polymer as a 2D self-avoiding polymer from synthesizing enzyme (only one shown for simplicity) exerts a radial outward areal pressure on the edge of the cell plate. The force acts within the plate only and thus includes cross-sectional area (not fenestrations).

## References

1. Cosgrove, D. J. 2005. Growth of the plant cell wall. Nature Reviews Molecular Cell Biology 6(11): 850–861.

2. Grones, P., S. Raggi, and S. Robert. 2019. FORCE-ing the shape. Current Opinion in Plant Biology 52:1–6.

3. Samuels, A. L., T. H. Giddings, and L. A. Staehelin. 1995. CYTOKINESIS IN TOBACCO BY-2 AND ROOT-TIP CELLS - A NEW MODEL OF CELL PLATE FORMATION IN HIGHER-PLANTS. Journal of Cell Biology 130(6): 1345–1357.

4. Segui-Simarro, J. M., J. R. Austin, E. A. White, and L. A. Staehelin. 2004. Electron tomographic analysis of somatic cell plate formation in meristematic cells of arabidopsis preserved by high-pressure freezing. Plant Cell 16(4): 836–856.

5. Smertenko, A., F. Assaad, F. Baluska, M. Bezanilla, H. Buschmann, G. Drakakaki, M.-T. Hauser, M. Janson, Y. Mineyuki, I. Moore, S. Mueller, T. Murata, M. S. Otegui, E. Panteris, C. Rasmussen, A.-C. Schmit, J. Samaj, L. Samuels, L. A. Staehelin, D. Van Damme, G. Wasteneys, and V. Zarsky. 2017. Plant Cytokinesis: Terminology for Structures and Processes. Trends in Cell Biology 27(12): 885–894.

6. Lee, Y.-R. J., and B. Liu. 2013. The rise and fall of the phragmoplast microtubule array. Current Opinion in Plant Biology 16(6): 757–763.

7. Drakakaki, G. 2015. Polysaccharide deposition during cytokinesis: Challenges and future perspectives. Plant Science 236:177–184.

8. Schweitzer, Y., T. Shemesh, and M. M. Kozlov. 2015. A Model for Shaping Membrane Sheets by Protein Scaffolds. Biophysical Journal 109(3): 564–573.

9. Shemesh, T., R. W. Klemm, F. B. Romano, S. Wang, J. Vaughan, X. Zhuang, H. Tukachinsky, M. M. Kozlov, and T. A. Rapoport. 2014. A model for the generation and interconversion of ER morphologies. Proceedings of the National Academy of Sciences of the United States of America 111(49): E5243–E5251.

10. Terasaki, M., T. Shemesh, N. Kasthuri, R. W. Klemm, R. Schalek, K. J. Hayworth, A. R. Hand, M. Yankova, G. Huber, J. W. Lichtman, T. A. Rapoport, and M. M. Kozlov. 2013. Stacked Endoplasmic Reticulum Sheets Are Connected by Helicoidal Membrane Motifs. Cell 154(2): 285–296.

11. Thiele, K., G. Wanner, V. Kindzierski, G. Jurgens, U. Mayer, F. Pachl, and F. F. Assaad. 2009. The timely deposition of callose is essential for cytokinesis in Arabidopsis. Plant Journal 58(1): 13–26.

12. Park, E., S. M. Diaz-Moreno, D. J. Davis, T. E. Wilkop, V. Bulone, and G. Drakakaki. 2014. Endosidin 7 Specifically Arrests Late Cytokinesis and Inhibits Callose Biosynthesis, Revealing Distinct Trafficking Events during Cell Plate Maturation. Plant Physiology 165(3): 1019–1034.

13. Chen, X.-Y., L. Liu, E. Lee, X. Han, Y. Rim, H. Chu, S.-W. Kim, F. Sack, and J.-Y. Kim. 2009. The Arabidopsis Callose Synthase Gene GSL8 Is Required for Cytokinesis and Cell Patterning. Plant Physiology 150(1): 105–113.

14. Guseman, J. M., J. S. Lee, N. L. Bogenschutz, K. M. Peterson, R. E. Virata, B. Xie, M. M. Kanaoka, Z. L. Hong, and K. U. Torii. 2010. Dysregulation of cell-to-cell connectivity and stomatal patterning by loss-of-function mutation in Arabidopsis CHORUS (GLUCAN SYNTHASE-LIKE 8). Development 137(10): 1731–1741.

15. Rawicz, W., K. C. Olbrich, T. McIntosh, D. Needham, and E. Evans. 2000. Effect of chain length and unsaturation on elasticity of lipid bilayers. Biophysical Journal 79(1): 328–339.

16. Lazaro, G. R., I. Pagonabarraga, and A. Hernandez-Machado. 2015. Phase-field theories for mathematical modeling of biological membranes. Chemistry and Physics of Lipids 185:46–60.

17. Choksi, R., M. Morandotti, and M. Veneroni. 2013. GLOBAL MINIMIZERS FOR AXISYMMETRIC MULTIPHASE MEMBRANES. Esaim-Control Optimisation and Calculus of Variations 19(4): 1014–1029.

18. Sarasij, R. C., S. Mayor, and M. Rao. 2007. Chirality-induced budding: A raft-mediated mechanism for endocytosis and morphology of caveolae? Biophysical Journal 92(9): 3140–3158.

19. Helfrich, W. 1973. ELASTIC PROPERTIES OF LIPID BILAYERS - THEORY AND POSSIBLE EXPERIMENTS. Zeitschrift Fur Naturforschung C-a Journal of Biosciences C 28(11-1):693–703.

20. Brakke, K. A. 1996. The surface evolver and the stability of liquid surfaces. Philosophical Transactions of the Royal Society a-Mathematical Physical and Engineering Sciences 354(1715): 2143–2157.

21. Lee, J. M. 1997. Riemannian Manifolds. Springer-Verlag New York, Inc.

22. Dimova, R. 2014. Recent developments in the field of bending rigidity measurements on membranes. Advances in Colloid and Interface Science 208:225–234.

23. Hu, M., J. J. Briguglio, and M. Deserno. 2012. Determining the Gaussian Curvature Modulus of Lipid Membranes in Simulations. Biophysical Journal 102(6): 1403–1410.

24. Fischer-Friedrich, E., A. A. Hyman, F. Juelicher, D. J. Mueller, and J. Helenius. 2014. Quantification of surface tension and internal pressure generated by single mitotic cells. Scientific Reports 4.

25. Kent, I. A., and T. P. Lele. 2017. Microtubule-based force generation. Wiley Interdisciplinary Reviews-Nanomedicine and Nanobiotechnology 9(3).

26. Schulmann, N., H. Meyer, T. Kreer, A. Cavallo, A. Johner, J. Baschnagel, and J. P. Wittmer. 2013. Strictly Two-Dimensional Self-Avoiding Walks: Density Crossover Scaling. Polymer Science Series C 55(1): 181–211.

27. Babb, V. M., and C. H. Haigler. 2001. Sucrose phosphate synthase activity rises in correlation with high-rate cellulose synthesis in three heterotrophic systems. Plant Physiology 127(3): 1234–1242.

28. Jung, H. T., B. Coldren, J. A. Zasadzinski, D. J. Iampietro, and E. W. Kaler. 2001. The origins of stability of spontaneous vesicles. Proceedings of the National Academy of Sciences of the United States of America 98(4): 1353–1357.

29. McMahon, H. T., and E. Boucrot. 2015. Membrane curvature at a glance. Journal of Cell Science 128(6): 1065–1070.

30. Ahn, G., H. Kim, D. H. Kim, H. Hanh, Y. Yoon, I. Singaram, K. J. Wijesinghe, K. A. Johnson, X. H. Zhuang, Z. Z. Liang, R. V. Stahelin, L. W. Jiang, W. Cho, B. H. Kang, and I. Hwang. 2017. SH3 Domain-Containing Protein 2 Plays a Crucial Role at the Step of Membrane Tubulation during Cell Plate Formation. Plant Cell 29(6): 1388–1405.

31. McMichael, C. M., and S. Y. Bednarek. 2013. Cytoskeletal and membrane dynamics during higher plant cytokinesis. New Phytologist 197(4): 1039–1057.

32. Otegui, M. S., D. N. Mastronarde, B. H. Kang, S. Y. Bednarek, and L. A. Staehelin. 2001. Three-dimensional analysis of syncytial-type cell plates during endosperm cellularization visualized by high resolution electron tomography. Plant Cell 13(9): 2033–2051.

33. Lou, H.-Y., W. Zhao, X. Li, L. Duan, A. Powers, M. Akamatsu, F. Santoro, A. F. McGuire, Y. Cui, D. G. Drubin, and B. Cui. 2019. Membrane curvature underlies actin reorganization in response to nanoscale surface topography. Proceedings of the National Academy of Sciences of the United States of America 116(46): 23143–23151.

34. Him, J. L. K., L. Pelosi, H. Chanzy, J. L. Putaux, and B. Bulone. 2001. Biosynthesis of (1 -> 3)-beta-D-glucan (callose) by detergent extracts of a microsomal fraction from Arabidopsis thaliana. (vol 268, pg 4628, 2001). European Journal of Biochemistry 268(21): 5653–5653.

35. Gidley, M. J., and K. Nishinari. 2009. Physico-chemistry of (1,3)-beta-Glucans. Chemistry, Biochemistry, and Biology of (1-->3)-Beta-Glucans and Related Polysaccharides:47–118.

36. Falch, B. H., and B. T. Stokke. 2001. Structural stability of (1 -> 3)-beta-D-glucan macrocycles. Carbohydrate Polymers 44(2): 113–121.

37. Stone, B. A., and A. E. Clarke. 1992. Chemistry and biology of 1,3-β-Glucans. La Trobe University Press.

38. Abou-Saleh, R. H., M. C. Hernandez-Gomez, S. Amsbury, C. Paniagua, M. Bourdon, S. Miyashima, Y. Helariutta, M. Fuller, T. Budtova, S. D. Connell, M. E. Ries, and Y. Benitez-Alfonso. 2018. Interactions between callose and cellulose revealed through the analysis of biopolymer mixtures. Nature Communications 9.

## Supplementary Information References

1. Abbena, E., S. Salamon, and A. Gray. 2017. Modern Differential Geometry of Curves and Surfaces with Mathematica. CRC Press.

2. Dimova, R. 2014. Recent developments in the field of bending rigidity measurements on membranes. Advances in Colloid and Interface Science 208:225–234.

3. Schulmann, N., H. Meyer, T. Kreer, A. Cavallo, A. Johner, J. Baschnagel, and J. P. Wittmer. 2013. Strictly Two-Dimensional Self-Avoiding Walks: Density Crossover Scaling. Polymer Science Series C 55(1): 181–211.

5. Samuels, A. L., T. H. Giddings, and L. A. Staehelin. 1995. CYTOKINESIS IN TOBACCO BY-2 AND ROOT-TIP CELLS - A NEW MODEL OF CELL PLATE FORMATION IN HIGHER-PLANTS. Journal of Cell Biology 130(6): 1345–1357.

